# Size-dependent susceptibility of lake phytoplankton to light stress: An implication for succession of large green algae in a deep oligotrophic lake

**DOI:** 10.1101/2021.07.15.452429

**Authors:** Takehiro Kazama, Kazuhide Hayakawa, Takamaru Nagata, Koichi Shimotori, Akio Imai, Kazuhiro Komatsu

## Abstract

Field observations of the population dynamics and measurements of photophysiology in Lake Biwa were conducted by size class (< vs. > 30 μm) from early summer to autumn to investigate the relationships between susceptibility to light stress and cell size. Also, a nutrient bioassay was conducted to clarify whether the growth rate and photosystem II (PSII) photochemistry of small and large phytoplankton are limited by nutrient availability. Large phytoplankton, which have lower intracellular Chl-*a* concentrations, had higher maximum PSII photochemical efficiency (*F_v_/F_m_*) but lower non-photochemical quenching (*NPQ_NSV_*) than small phytoplankton under both dark and increased light conditions. The nutrient bioassay revealed that the PSII photochemistry of small phytoplankton was restricted by N and P deficiency at the pelagic site even at the end of the stratification period, while that of large phytoplankton was not. These results suggest that large phytoplankton have lower susceptibility to PSII photodamage than small phytoplankton due to lower intracellular Chl-*a* concentrations. The size dependency of susceptibility to PSII photoinactivation may play a key role in large algal blooms in oligotrophic water.

## Introduction

Global warming due to recent climate change is thought to cause downward selection pressure on the cell size of phytoplankton through changes to the thermal stratification of lakes and oceans (Finkel et al., 2009, 2010; Winder et al., 2009). In lakes, long-term warming and seasonal increases in water temperature intensify the strength of stratification and promote oligotrophication of the offshore surface layer (Zohary et al., 2020). Generally, smaller phytoplankton cells have advantages in the upper layer of stratified water columns (Falkowski & Oliver, 2007; Winder et al., 2009; Zohary et al., 2020) because of a lower sinking velocity (Reynolds, 2006), rapid growth rate (Banse, 1976; Schlesinger et al., 1981; Finkel et al., 2010; Mei et al., 2011) and a relatively high nutrient uptake rate per cell volume (Suttle et al., 1987; Sunda & Hardison, 2010; Edwards et al., 2011). Conversely, larger phytoplankton cells are generally associated with eutrophic environments of well-mixed water columns (Sommer et al., 1986, 2012).

In addition to these factors, reduced vertical mixing also changes the light environment, which can play a determining role in the size of phytoplankton cells (Cermeño et al., 2005; Finkel et al., 2009; Key et al., 2010; Rugema et al., 2019). Jin et al. (2013) pointed out that strengthened stratification can increase light intensity and reduce light fluctuation, which may influence photosynthetic activity and productivity of phytoplankton in upper mixed layers. Recent bio-optical studies suggest that species composed of relatively large cells may be less susceptible to excess light or fluctuation stress than those composed of smaller cells (Suggett et al., 2009; Alderkamp et al., 2010; Key et al., 2010). Species composed of large cells and those that form large aggregated colonies tend to have lower intracellular Chl-*a* concentrations (C_i_) to avoid intracellular self-shading effects (i.e. the packaging effect, Duyens, 1956; Agustí, 1991; Finkel, 2001). Low C_i_, leads to decreased light absorption cross sections and, subsequently, lower sensitivity to photosystem II (PSII) photoinactivation (Finkel et al., 2010; Key et al., 2010) due to excess light energy (Marshall et al., 2000). Photoinactivation not only reduces linear electron flow in PSII but also increases the energy required for PSII repair (Key et al., 2010; Campbell & Serôdio, 2020). Thus, high light conditions can be particularly advantageous to large phytoplankton (Key et al., 2010). However, relatively few studies of marine diatoms and picocyanobacteria have focused on the size dependence of population dynamics and photophysiology in natural communities (Suggett et al., 2009; Key et al., 2010; Giannini & Ciotti, 2016).

Lake Biwa is a deep oligotrophic lake in Japan with well documented physical and chemical conditions and phytoplankton community compositions (Kawanabe et al., 2012). According to a previous study, the mean size of phytoplankton cells had been decreasing along with increasing water temperature until 2009 after a bloom of *Staurastrum dorsidentiferum* var. *ornatum* Glönblad 1983 (hereafter, *S. dorsdeintiferum*), a large member of the class Zygnematophyceae, in the 1980s (Kishimoto et al., 2013). However, a larger Zygnematophyceae species, *Micrasterias hardyi* West, successfully invaded in the early 2000s and dominated the North Basin during the winter of 2016 (Hodoki et al., 2020). This counter-intuitive phenomenon was not associated with nutrient loading, as total phosphorus had been maintained at an exceptionally low concentration of < 0.01 mg L^-1^ (N{Citation}ishino, 2012; Shiga Prefecture, 2018). Hence, a detailed investigation of the natural phytoplankton community by size is needed to clarify how populations of large-celled algae are established in an oligotrophic lake.

The working hypothesis of this study is that the large phytoplankton community in Lake Biwa has a low pigment concentration and, thus, low susceptibility to light stress due to low C_i_. Field observations of the population dynamics and photophysiology measurement by size class (S, < 30 μm and L, > 30 μm) in Lake Biwa were conducted from early summer to autumn. Also, growth experiments with N and P enrichment were conducted to clarify whether the growth rate and PSII photochemistry of small and large phytoplankton are limited by nutrient availability. Multi-excitation wavelength fast repetition rate fluorometry (FRRf) was used to evaluate the photophysiology of the phytoplankton community. FRRf was conducted with the use of three different excitation lights corresponding to the absorption spectra of phycobilisomes in cyanobacteria (Kazama et al., 2021a). This nonintrusive bio-optical method enables assessment of PSII status of the algal community at different spatial and time scales by measuring the maximum photochemical efficiency (*F_v_/F_m_*; in a dark-adapted state), effective photochemical efficiency (*F_q_′/F_m_′*; in a light-adapted state) and normalized Stern-Volmer coefficient of quenching (*NPQ_NSV_*) (Kolber & Falkowski, 1993; Kolber et al., 1998; McKew et al., 2013, see Abbreviations). Increases in *F_v_/F_m_* and *F_q_′/F_m_′* values imply increasing active PSII sites by synthesis or repair (Cosgrove & Borowitzka, 2010), while an increase in the *NPQ_NSV_* value implies an increase in the ratio of total non-photochemical quenching to the photochemistry rate constant to alleviate excess excitation pressure (Mckew et al. 2013).

## Materials and Methods

### Sampling procedure

Sampling was conducted at two long-term survey stations established in the North Basin of Lake Biwa on Honshu Island, Japan (Fig. 1), station 12B (62 m depth, 35°11’39” N, 135°59’39” E) and station 12C (7 m depth, 35°10’40” N, 136°03’07” E), on 22 June, 10 July, 24 August, 7 September, 26 October and 23 November 2019. No permits were required to collect water samples from the lake. Vertical profiles of irradiance and water temperature were measured using a water quality sonde (AAQ-RINKO; JFE Advantech Co., Ltd., Nishinomiya, Japan). Water samples were collected into 5-L Niskin bottles on a rosette sampler (AWS; JFE Advantech Co. Ltd.) at a depth of 5 m at station 12B and into 10-L Niskin samplers at a depth of 2.5 m at station 12C. Samples were stored in plastic bags in the dark and transferred to the laboratory for chemical and biological analyses within 2 h after collection.

**Fig. 1.**
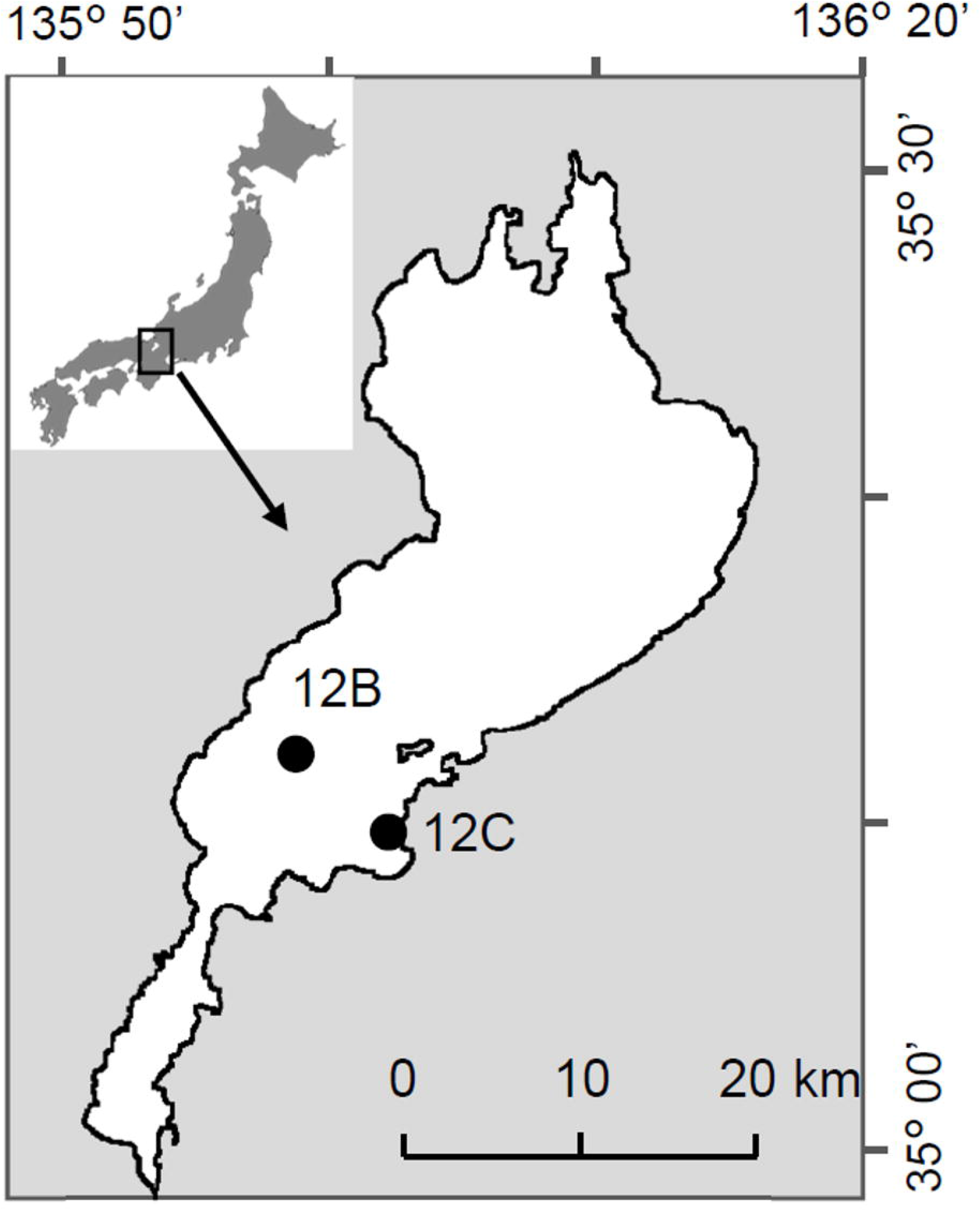
Map of the study sites in Lake Biwa, Japan This figure was reproduced from the website of the Geospatial Information Authority of Japan (https://www.gsi.go.jp) and supplemented with latitude and longitude lines. This map is licensed under the Government of Japan Standard Terms of Use (Ver. 2.0), which are compatible with the Creative Commons Attribution License 4.0 (CC BY 4.0).

Macro-nutrient concentrations were determined from a 100-mL aliquot of a subsample collected at each depth. The subsample was immediately filtered through a syringe-type membrane filter (0.2 μm pore size, Acrodisc syringe filter; Pall Corporation, Ann Arbor, MI, USA). The filtered samples were stored at −20°C until analyses of nitrate, ammonia and phosphate concentrations using an ion chromatograph system (Dionex Integrion HPIC system; Thermo Scientific, Waltham, MA, USA). For chlorophyll a (Chl-a) analysis, 50-300-mL samples were preliminary filtered through 200-μm nylon mesh to remove crustacean zooplankton (major grazers in Lake Biwa) and then separated into small (S, < 30 μm) and large (L, > 30 μm) fractions using nylon mesh. For the small size Chl-*a* fraction, samples were passed through a 30-μm nylon mesh and then filtered with a 25-mm glass fiber filter (0.7 μm nominal pore size, GF/F; GE Healthcare, UK Inc., Little Chalfont, UK). For the large size Chl-*a* fraction, samples remaining on the 30-μm nylon mesh were washed with Milli-Q water and filtered through a GF/F filter. Chl-*a* was extracted with N,N-dimethylformamide for 24 h in the dark (Suzuki & Ishimaru, 1990) and then stored at −80°C. The Chl-*a* concentration was determined with a 10-AU fluorometer (Turner Designs, Sunnyvale, CA, USA). To determine the number of phytoplankton, a 50-mL aliquot of each sample was fixed with Lugol’s solution (1% final concentration). After 24 h of settling in the dark, the supernatant was gently removed and the sample was concentrated to 15 mL. All phytoplankton cells were enumerated at the finest taxon level (species or genus) under a light microscope at ×100-400 magnification at the Marine Biological Research Institute of Japan (Tokyo, Japan). The species with large variations in cell size were separated into several size classes. Cell density was calculated from the cell or colony density and the average cell number of the colony. For estimation of cell volume (V_cell_), the dimensions of 10 to 20 cells for each taxon were measured based on the work of Hillebrand et al. (1999) and Olenina et al. (2006). Selected cell shapes for all taxa are shown in Appendix Table 1. The phytoplankton community composition was assessed based on the carbon biomass converted from the V_cell_ (Menden-Deuer & Lessard, 2000). Chl-*a* concentration per volume (C_i_, pg μm^-3^) was calculated for each size class.

### Photophysiology of small and large phytoplankton

Preliminary screened samples were fractionated into two size classes as mentioned above. Large phytoplankton were resuspended in lake water filtered through a GF/F filter and diluted to the *in situ* concentration with filtered lake water. For photophysiology measurements, a 2-mL aliquot of each size class was poured into two 10-mL glass tubes. Samples were acclimated in the dark for 20 min at the *in situ* temperature ± 1 °C in a growth chamber (HCLP-880PF; Nippon Medical and Chemical Instruments Co., Ltd., Osaka, Japan). Photophysiological characteristics of dark-acclimated samples were then measured with a bench-top FRRf (Act2; Chelsea Technologies Ltd., West Molesey, UK) equipped with three light-emitting diodes (LEDs) that provide flash excitation energy centred at 444, 512 and 633 nm (Kazama et al., 2021a). Here, 444 nm (blue) corresponds to the absorption peak of Chl-*a*, while 512 nm (green) and 633 nm (orange) correspond to the absorption peaks of phycoerythrin and phycocyanin (Wojtasiewicz & Stoń-Egiert, 2016). A combination of all three LEDs was employed to precisely measure the minimum PSII fluorescence yield (*F_O_*) of the natural communities, including cyanobacteria (Kazama et al., 2021a). A single turnover method was applied that consisted of a saturation phase (100 flashlets with a 2-μs pitch) and a relaxation phase (40 flashlets with a 60-μs pitch). This sequence was repeated 16 times with a 200-ms interval between acquisitions. Measurement was repeated five times per sample. The power of the flashlets and the gain of the extra high tension of the photomultiplier tube (PMT eht) were optimised with Act2Run software (version 2.4.1.0; Chelsea Technologies, Ltd.). After 60-s measurements in the dark, the samples were immediately exposed to 40-s periods of 5 to 8 actinic lights increasing stepwise in intensity from 0 to 850 μmol photon m^-2^ s^-1^. Baseline fluorescence was determined from the sample passed through a 0.2-μm pore-size Acrodisc syringe filters and subtracted from the total variable fluorescence signal, as described by Hughes et al. (2018). The *F_v_/F_m_* and *F_q_′/F_m_′* values under actinic light were evaluated as the efficiency of PSII photochemistry and *NPQ_NSV_* (*F_O_′/F_v_′*) as non-photochemical quenching. Quality control of all FRRf data was assessed as described in a previous study (Kazama et al., 2021a). In short, *RσPSII* (the probability of an RCII being closed during the first flashlet of a single turnover saturation phase under dark) or *RσPSII′* (same to *RσPSII* but under actinic light) values of < 0.03 or > 0.08 were rejected.

### Nutrient bioassay

To assess the dependency of cell size, growth rate and PSII photochemistry on nutrient limitations in phytoplankton, N and P enrichment experiments were conducted. In brief, 400 mL of lake water (preliminary screened with 200-μm nylon mesh) were poured into six 500-mL polycarbonate bottles (Nalgene, Rochester, NY, USA). Three bottles were spiked with NaNO_3_ and K_2_HPO_4_ to a final concentration of 32 and 2 μM (NP), respectively, while the other bottles received no nutrients and served as controls (Con). All bottles were incubated for 48 h under 500 μmol photon m^-2^ s^-1^ at the *in situ* temperature ± 1°C in a growth chamber (HCLP-880PF). The nutrient concentrations and incubation times were chosen to ensure stimulation of phytoplankton growth as described by Kagami & Urabe (2001). Light was provided under a 12:12-h light:dark cycle with a 20 W orange LED (PF20-S9WT8-D; Nippon Medical and Chemical Instruments Co., Ltd.).

After incubation, the Chl-*a* concentration, phytoplankton biomass, *F_v_/F_m_* ratio and *NPQ_NSV_* value of the dark-adapted samples were measured as mentioned above. The apparent population growth rate of Chl-*a* (μ[chl], d^-1^) and biomass (μ[mass], d^-1^) were calculated assuming exponential growth as follows:

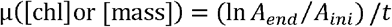

where *A_ini_* and *A_end_* are the Chl-*a* concentrations or biomass of each size class at the beginning and the end of the incubation, respectively, and *t* is incubation time.

### Statistical analyses

Potential correlations of any two factors among the V_cell_, C_i_, *F_v_/F_m_* and *NPQ_NSV_* values were identified using the Spearman’s rank correlation coefficient (*ρ*) with a significance level of *p* < 0.05. The effects of nutrient enrichment and size fraction on μ[chl], μ[mass], C_i_, *F_v_/F_m_* and *NPQ_NSV_* were examined by two-way analysis of variance (ANOVA) followed Tukey-Kramer’s post-hoc test with a significance level of *p* < 0.05 using the aov() and TukeyHSD() functions of R software version 3.6.3 (https://www.r-project.org/; R Development Core Team, 2020). The effect of nutrient enrichment and size fraction on phytoplankton species composition was examined by permutational multivariate ANOVA (PERMANOVA) with a significance level of *p* < 0.05 using the adonis() function in the R package ‘vegan’ (Oksanen et al., 2018) with the package ‘MASS’ (Ripley et al., 2020).

## Results

### Environments

Ancillary measurements of water temperature, light and concentrations of NO_3_^-^, NH_4_^+^ and PO_4_^3+^ of each sample showed clear spatial and seasonal variability (Table 1, Fig. 2). Water temperature varied from 16.3°C to 29.0°C and from 14.8°C to 28.8°C at stations 12B and 12C, respectively, during the study period. At station 12B, the thermocline was shallower than the euphotic zone depth in July, August and September. At the end of the stratification period, the depth of the thermocline was 20 m or more in October and November. At station 12C, although stratification was weaker than that at station 12B, the water temperature decreased gradually with depth from 30°C to 26°C in August. Light penetrated to the bottom layer at station 12C during the study period. The subsurface Chl-*a* fluorescence maximum was observed from June to September at station 12B (Fig. 2). At station 12C, Chl-*a* fluorescence peaked in the mid or bottom layer.

**Table 1.**
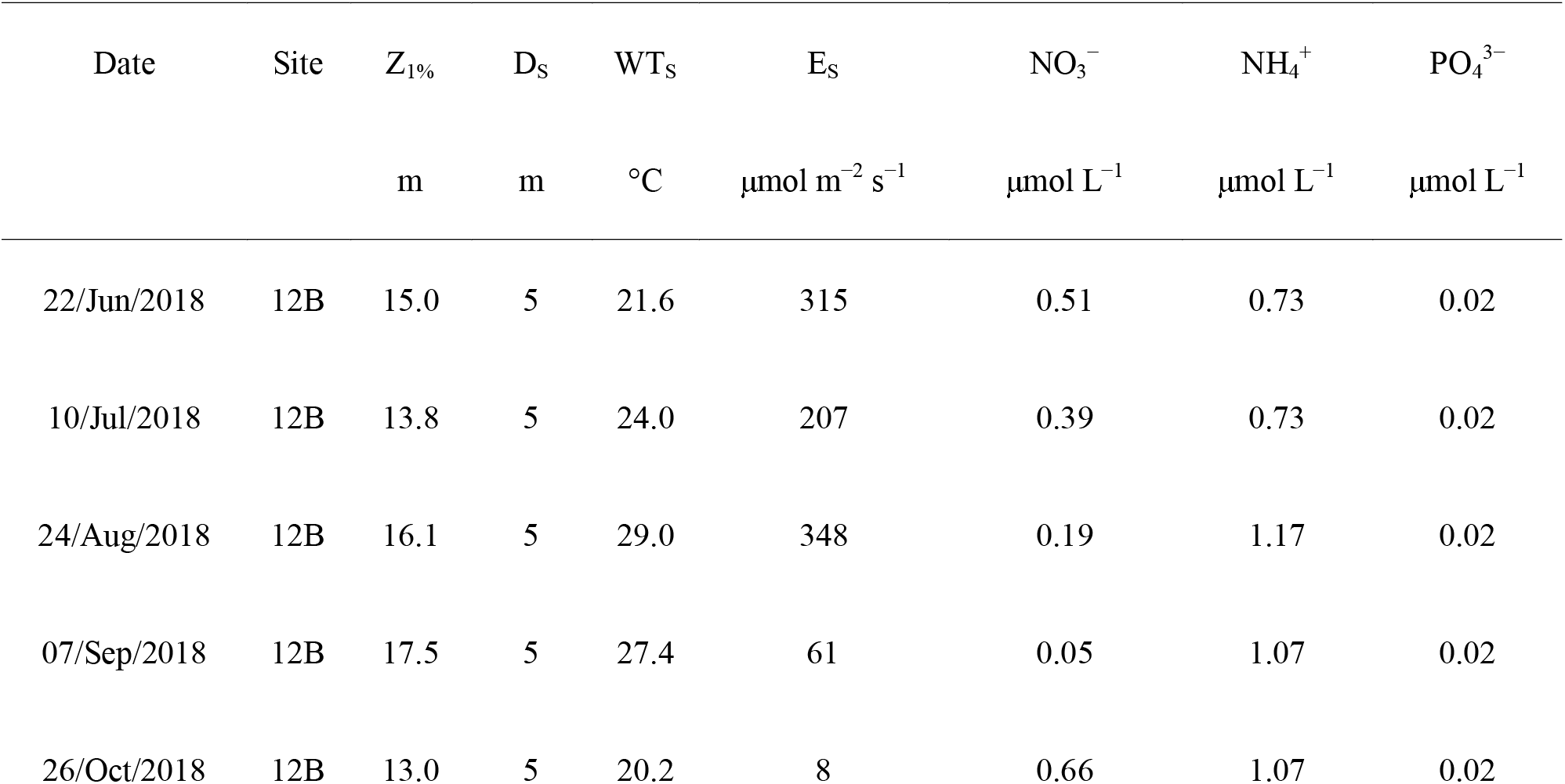
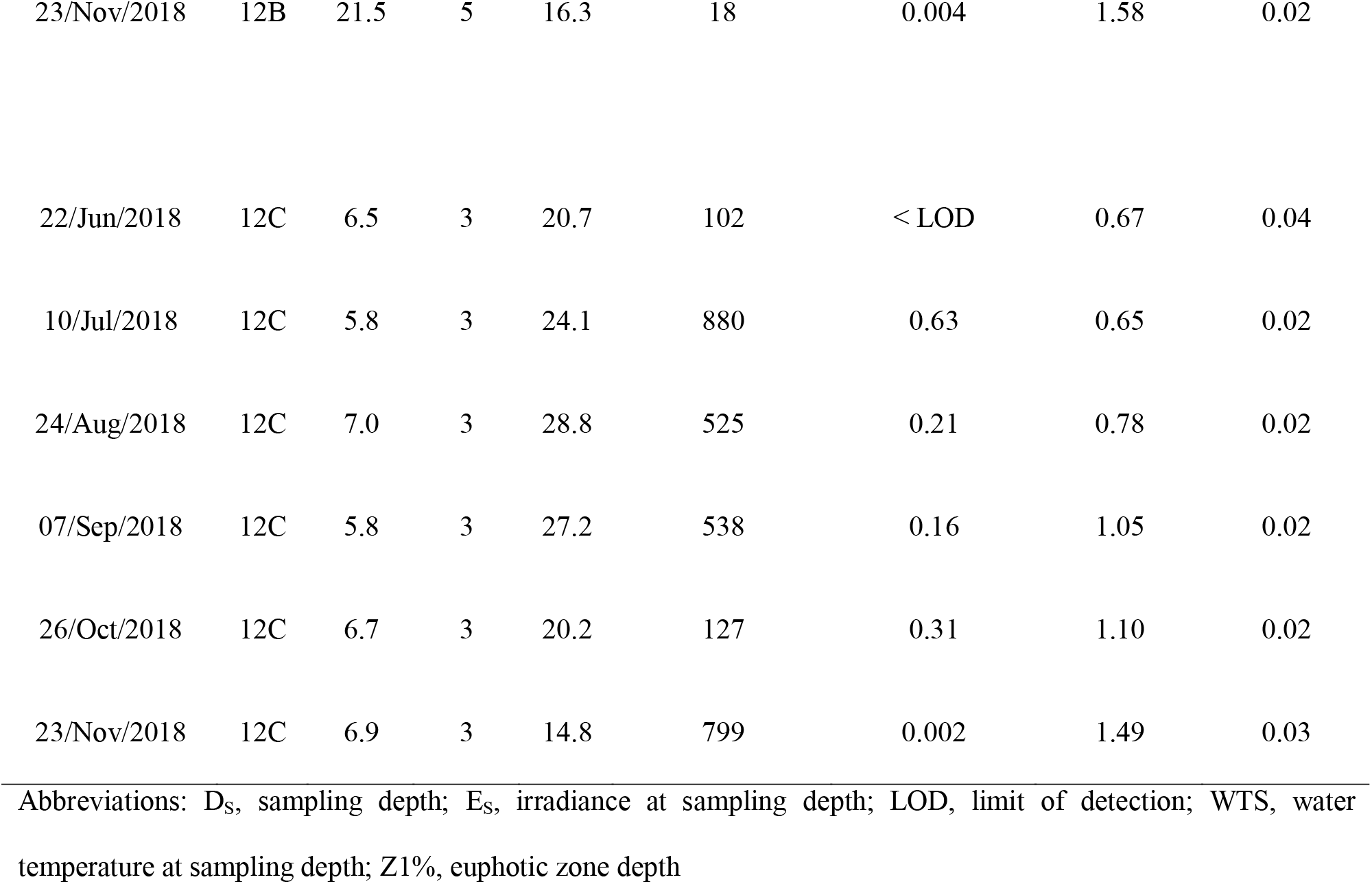
Physical and chemical conditions at the sampling stations.

| Date | Site | $Z_{1\%}$ | $D_s$ | $WT_s$ | $E_s$ | $\text{NO}_3^-$ | $\text{NH}_4^+$ | $\text{PO}_4^{3-}$ |
| --- | --- | --- | --- | --- | --- | --- | --- | --- |
| | | m | m | °C | $\mu\text{mol m}^{-2} \text{s}^{-1}$ | $\mu\text{mol L}^{-1}$ | $\mu\text{mol L}^{-1}$ | $\mu\text{mol L}^{-1}$ |
| 22/Jun/2018 | 12B | 15.0 | 5 | 21.6 | 315 | 0.51 | 0.73 | 0.02 |
| 10/Jul/2018 | 12B | 13.8 | 5 | 24.0 | 207 | 0.39 | 0.73 | 0.02 |
| 24/Aug/2018 | 12B | 16.1 | 5 | 29.0 | 348 | 0.19 | 1.17 | 0.02 |
| 07/Sep/2018 | 12B | 17.5 | 5 | 27.4 | 61 | 0.05 | 1.07 | 0.02 |
| 26/Oct/2018 | 12B | 13.0 | 5 | 20.2 | 8 | 0.66 | 1.07 | 0.02 |
| 23/Nov/2018 | 12B | 21.5 | 5 | 16.3 | 18 | 0.004 | 1.58 | 0.02 |
| 22/Jun/2018 | 12C | 6.5 | 3 | 20.7 | 102 | < LOD | 0.67 | 0.04 |
| 10/Jul/2018 | 12C | 5.8 | 3 | 24.1 | 880 | 0.63 | 0.65 | 0.02 |
| 24/Aug/2018 | 12C | 7.0 | 3 | 28.8 | 525 | 0.21 | 0.78 | 0.02 |
| 07/Sep/2018 | 12C | 5.8 | 3 | 27.2 | 538 | 0.16 | 1.05 | 0.02 |
| 26/Oct/2018 | 12C | 6.7 | 3 | 20.2 | 127 | 0.31 | 1.10 | 0.02 |
| 23/Nov/2018 | 12C | 6.9 | 3 | 14.8 | 799 | 0.002 | 1.49 | 0.03 |

Abbreviations: D<sub>s</sub>, sampling depth; E<sub>s</sub>, irradiance at sampling depth; LOD, limit of detection; WTS, water temperature at sampling depth; Z1%, euphotic zone depth

**Fig. 2.**
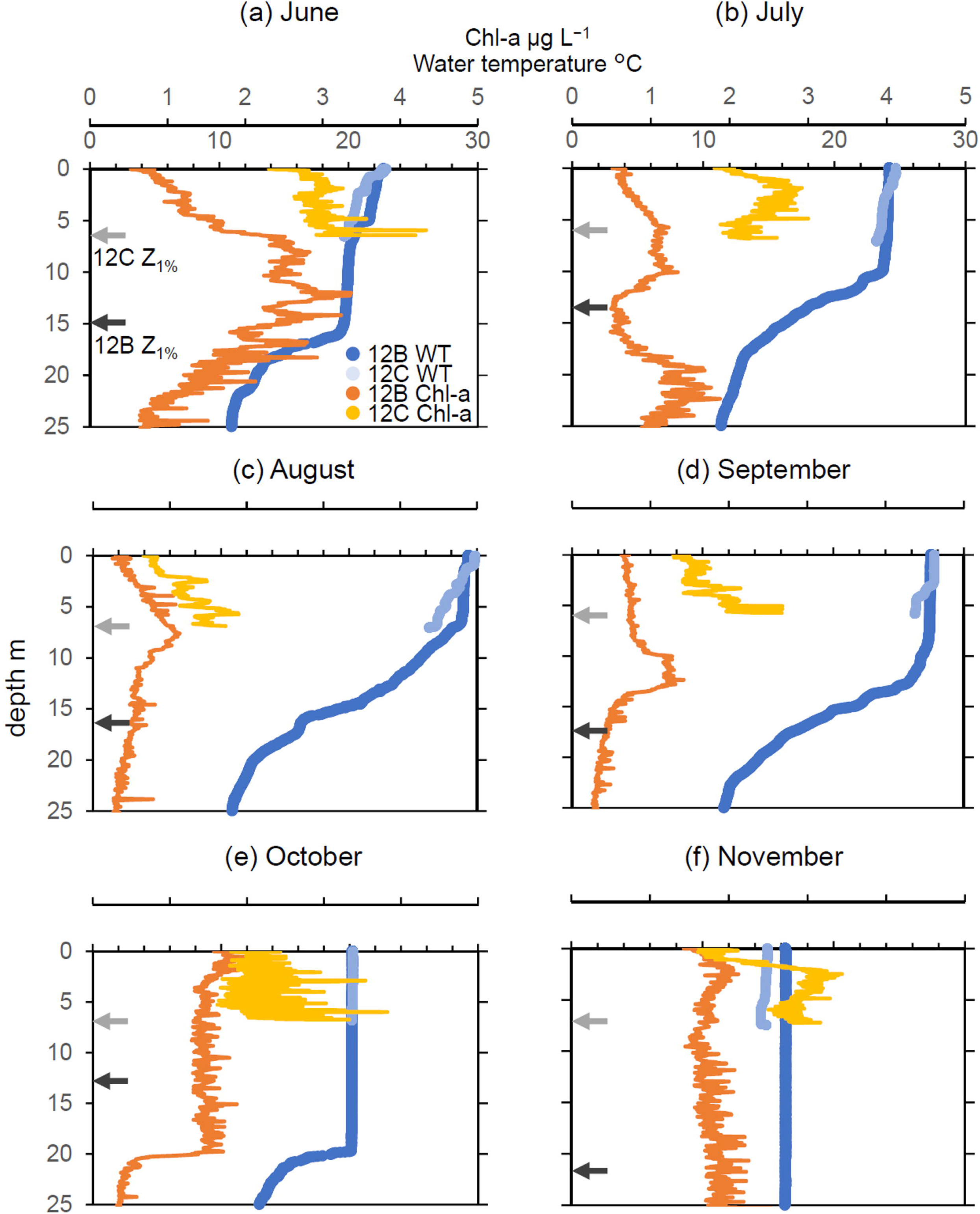
Vertical profiles of euphotic zone depth (Z_1%_), water temperature (WT) and Chl-*a* fluorescence of the upper 25 m at station 12B and 7 m at station 12C

Nutrient variation was similar at both stations. The NO_3_^-^ concentration was higher in June and October and the lowest in November, while the NH_4_^+^ concentration was lower in June and July and the highest in November. Meanwhile, the PO_4_^3+^ concentration remained at less than 0.02 μmol L^-1^ at station 12B and less than 0.04 μmol L^-1^ at station 12C throughout the study period.

### Phytoplankton abundance, morphological traits and species composition

By size fraction, the Chl-*a* concentration widely varied with space and time (Fig. 3a). The Chl-*a* concentration of the small size fraction at station 12B (12B-S) was around 2 μg L^-1^ throughout the study period, while that at station 12C (12C-S) peaked at 12 μg L^-1^ in June. The Chl-*a* concentration of the large size fraction at station 12B (12B-L) increased in October and November, while that at station 12C (12C-L) increased in July and October. There were clear differences in the biomass abundance between the small and large phytoplankton fractions (Fig. 3b). The biomass of the large phytoplankton fraction was always greater than that of the small phytoplankton fraction, with the exception of station 12B in July. In October and November, the biomasses of the small and large phytoplankton fractions increased at both stations. Similarly, the V_cell_ value at both stations increased by up to three-fold from September to October (Fig. 3c). The C_i_, of the small size fraction at stations 12B and 12C was always greater than that of the large size fraction throughout the study period (Fig. 3d).

**Fig. 3.**
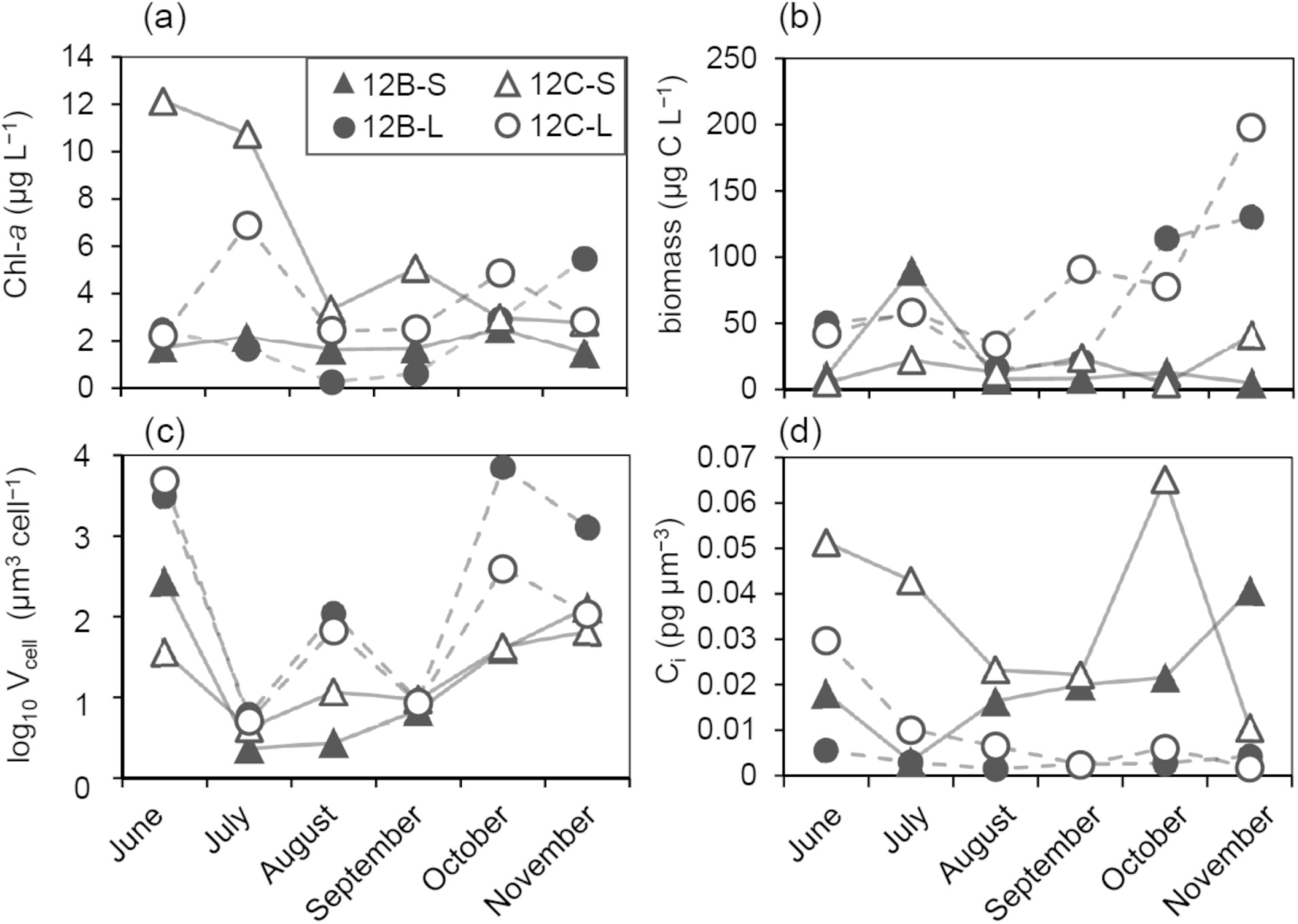
Seasonal variations in (a) Chl-*a* concentration, (b) carbon biomass, (c) V_cell_ and (d) C_i_, of the small (S) and large (L) phytoplankton fractions at stations 12B and 12C

In the small size fraction, cyanobacteria and diatoms dominated from June to October in both stations, while cryptophytes (station 12B) and small chlorophytes (station 12C) dominated in November (Figs. 4a, c). In the large size fraction, the abundance of large aggregated cyanobacterial colonies increased from July to September and that of zygnematophytes (mainly *S. dorsidentiferum* and *M. hardyi*) increased in June, October and November (Figs. 4b, d, Appendix Fig. A1). Chrysophytes, dinoflagellates and small flagellates always accounted for less than 20% of the total phytoplankton biomass.

**Fig. 4.**
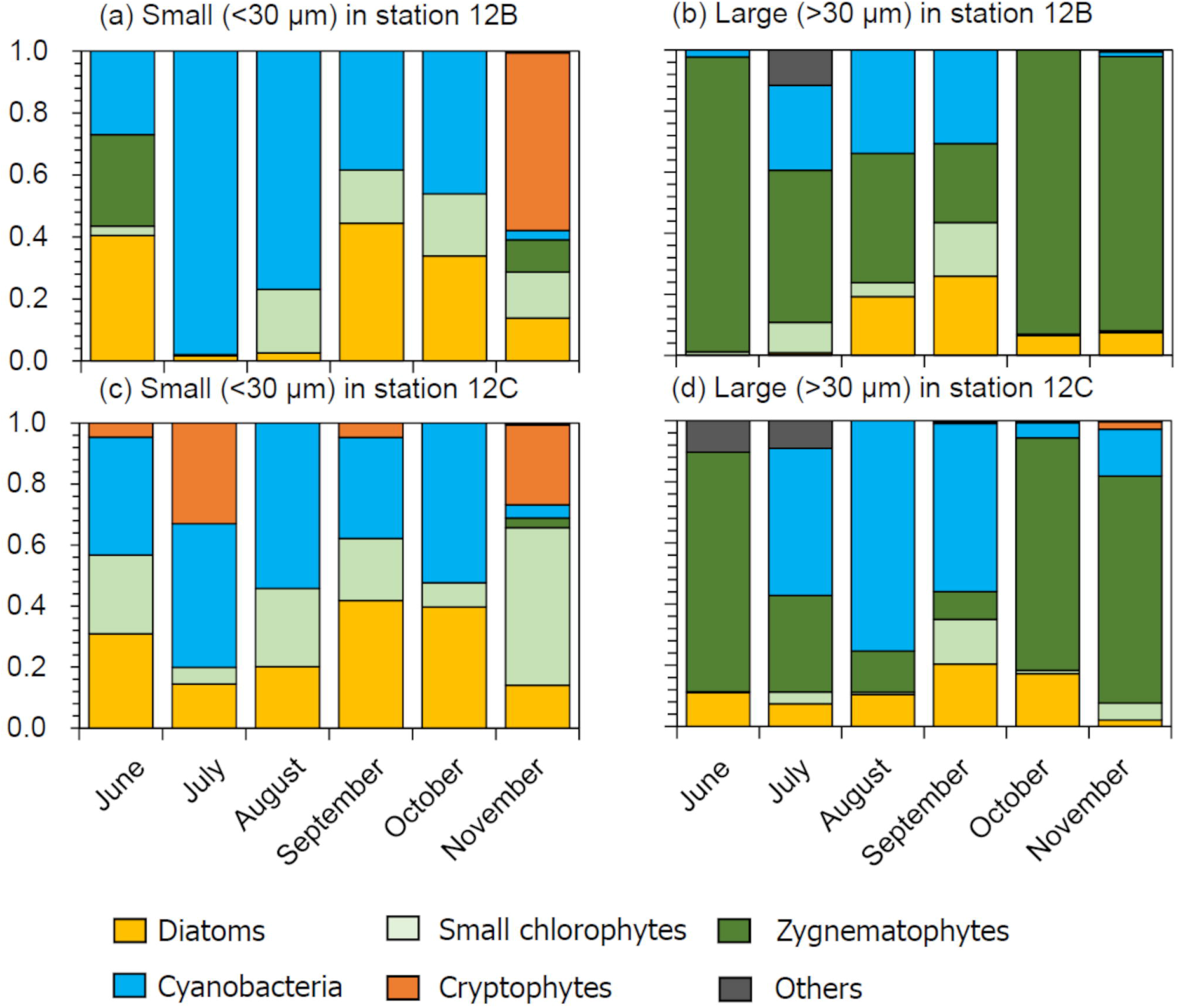
Monthly variation in phytoplankton composition based on carbon biomass of the small (a, c) and large (b, d) fractions at stations 12B (a, b) and 12C (c, d)

### PSII photophysiology

The *F_v_/F_m_* ratio was always greater in the large fraction than the small fraction at both stations, with the exception of station 12B in September (Fig. 5a). In contrast, the *NPQ_NSV_* value in the dark-adapted state was always greater in the small size fraction at both stations, with the exception of station 12B in September (Fig. 5b).

**Fig. 5.**
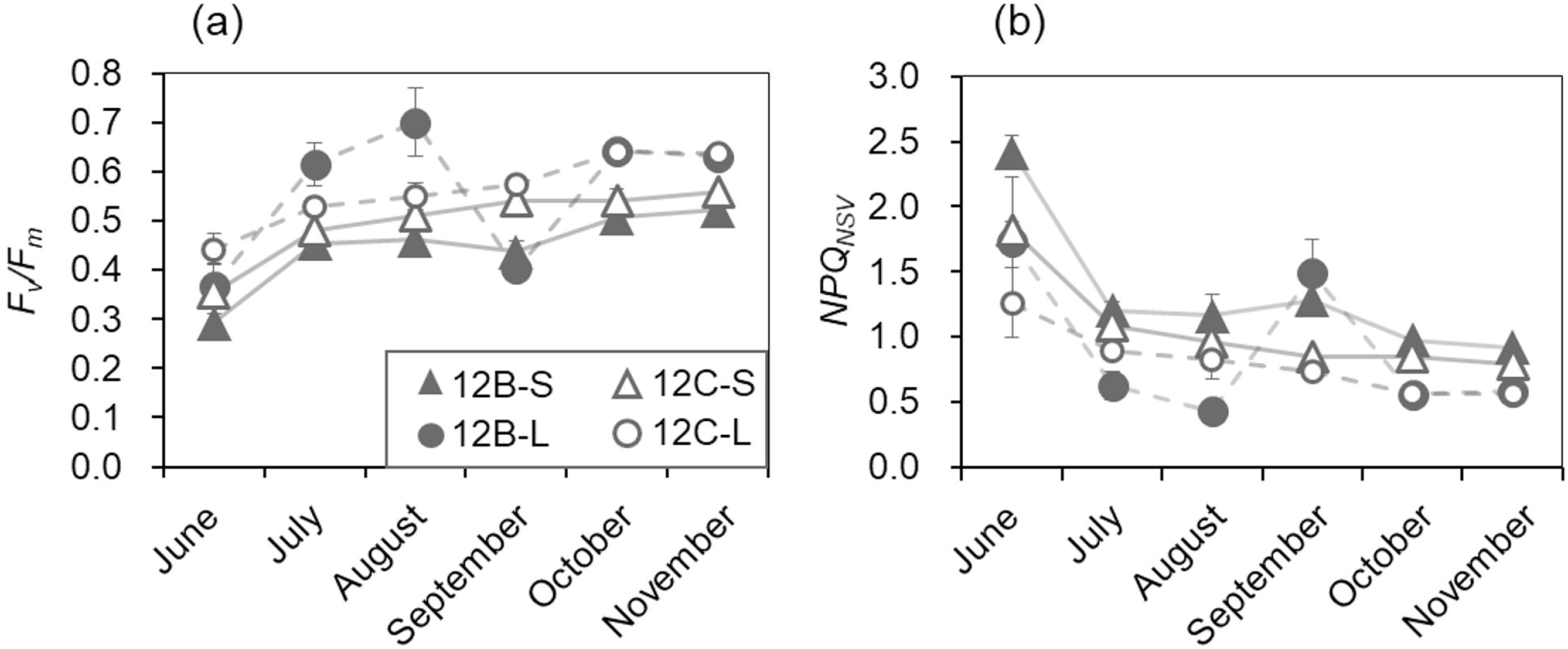
Variations in the *F_v_/F_m_* ratio (a) and *NPQ_NSV_* value (b) in the dark-adapted small and large phytoplankton fractions at each site Data are presented as mean values. Error bars denote the standard error (SE).

The *F_q_′/F_m_′* ratio of both size fractions decreased with increasing actinic light intensity (Fig. 6). However, the *F_q_′/F_m_′* ratio of the large size fractions was always greater than that of the small fractions with the exception of that in September at station 12B and June at station 12C. Conversely, the *NPQ_NSV_* value increased with increasing actinic light intensity (Fig. 7) and was always greater in the small fractions than the large fractions with the exception of that in September at station 12B and June at station 12C.

**Fig. 6.**
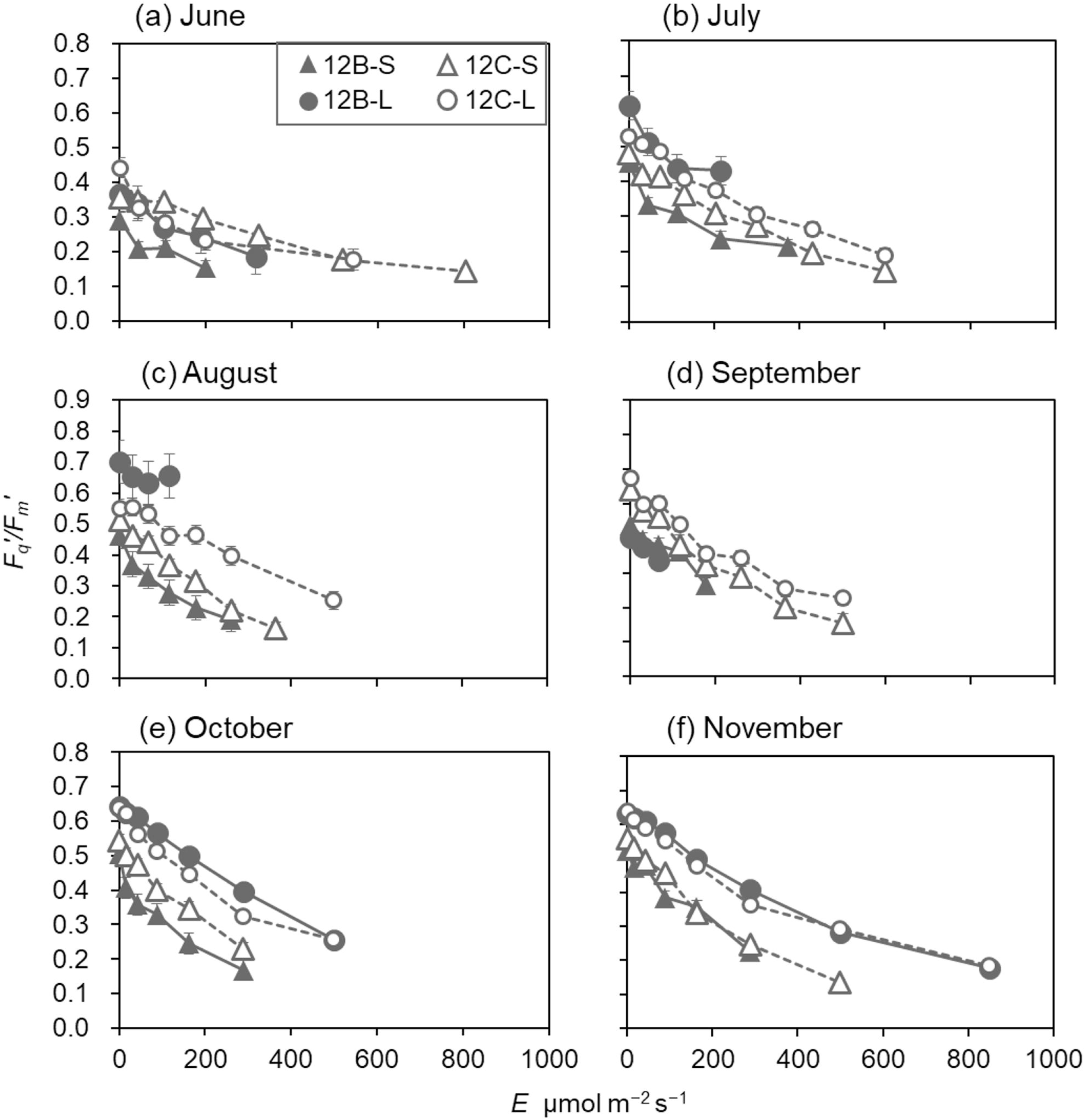
Responses of phytoplankton communities (*F_q_′/F_m_′*) to increasing light intensity at stations 12B and 12C Data are presented as mean values. Error bars denote the SE

**Fig. 7.**
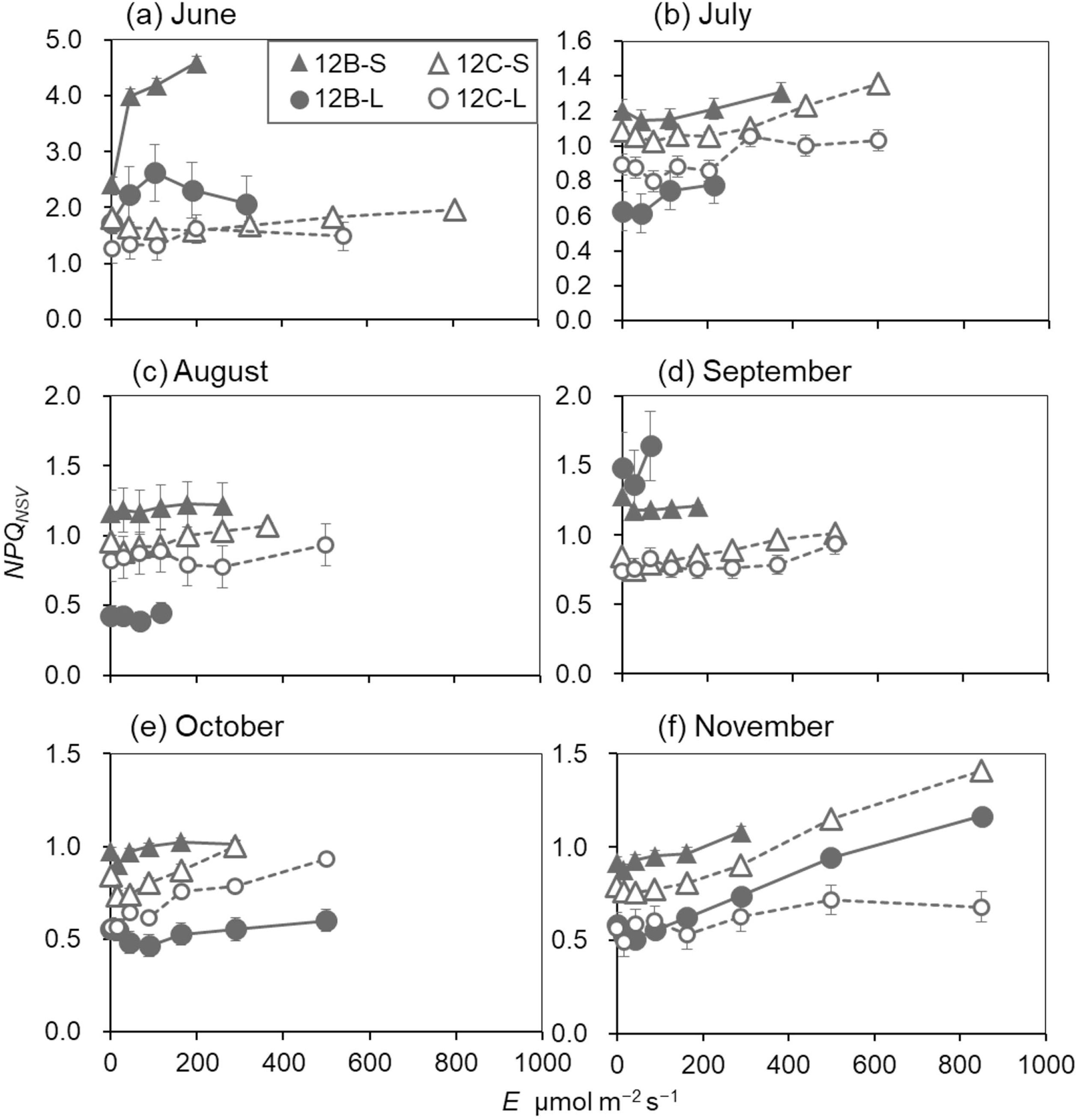
Responses of phytoplankton communities (*NPQ_NSV_*) to increasing light intensity at stations 12 B and 12C Data are presented as mean values. Error bars denote the SE.

The relationships between PSII photophysiology and cell size and C_i_, were also examined (Fig. 8). There was no correlation of the *F_v_/F_m_* ratio and *NPQ_NSV_* value with the V_cell_ value (*F_v_/F_m_*: ρ = 0.19, *p* = 0.37; *NPQ_NSV_*: ρ = −0.24, *p* = 0.25). However, the *F_v_/F_m_* ratio was negatively correlated with C_i_, (*ρ* = −0.45, *p* = 0.03), while the *NPQ_NSV_* was positively correlated with C_i_, (*ρ* = 0.49, *p* = 0.016).

**Fig. 8.**
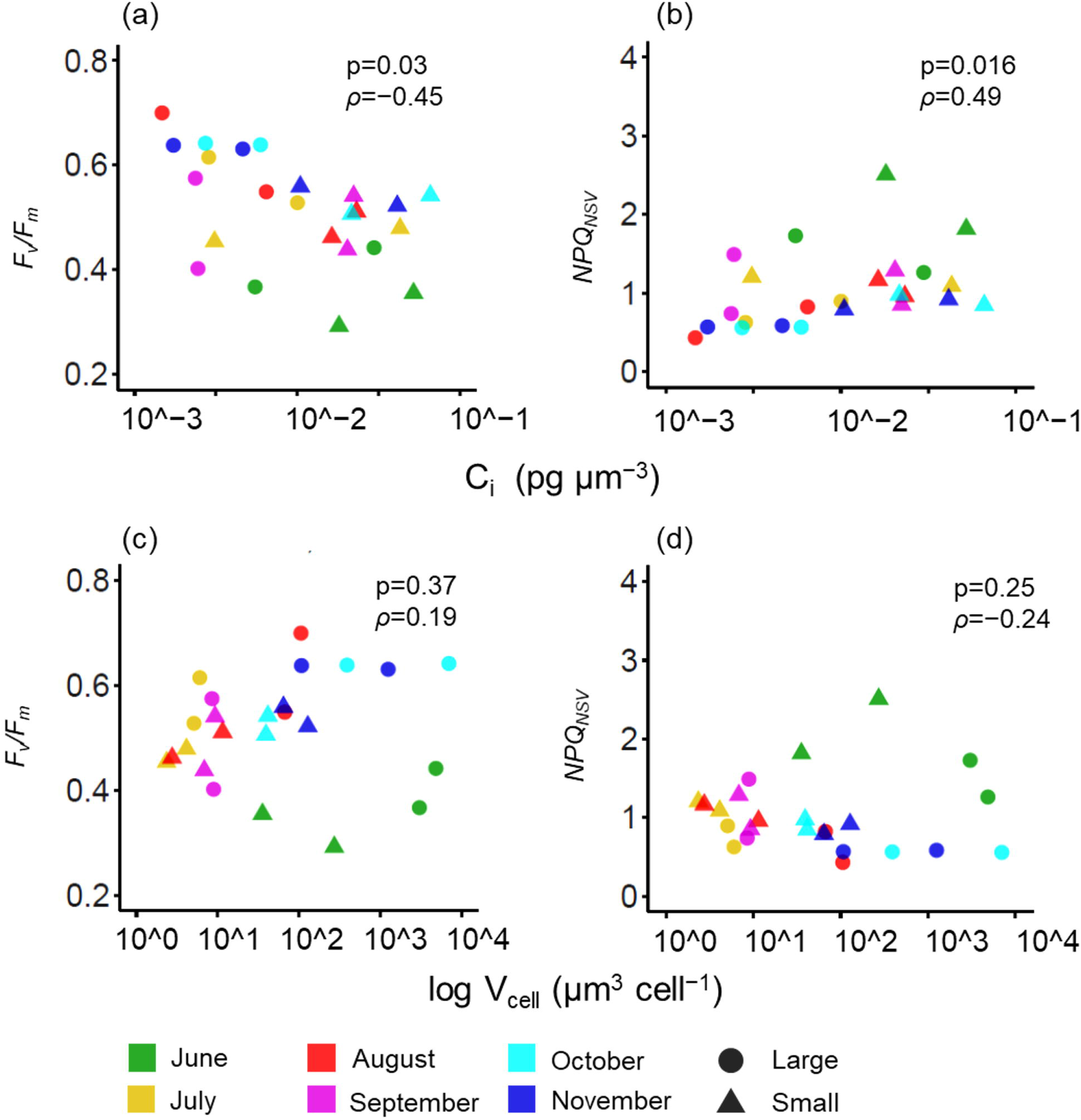
Scatter plots of C_i_, vs. the *F_v_/F_m_* ratio (a) and *NPQ_NSV_* value (b), and of the V_cell_ value vs. the *F_v_/F_m_* ratio (c) and *NPQ_NSV_* value (d) in dark-adapted small and large phytoplankton fractions at each site from June to November. Spearman’s *ρ* and *p* values are also shown

### Nutrient bioassay

The effects of N and P enrichment on the growth rate via the Chl-*a* concentration (μ[chl]), *F_v_/F_m_* ratio and *NPQ_NSV_* value of the small and large size phytoplankton fractions are summarized in Table 2 (bar plots are available in Appendix Fig. A2). In July, August and September, N and P enrichment significantly increased μ[chl] in both size fractions at both stations. After enrichment, there was no significant difference in μ[chl] between the size fractions at station 12B, while there were significant differences at station 12C (small > large in July and small < large in August and September).

**Table 2.**
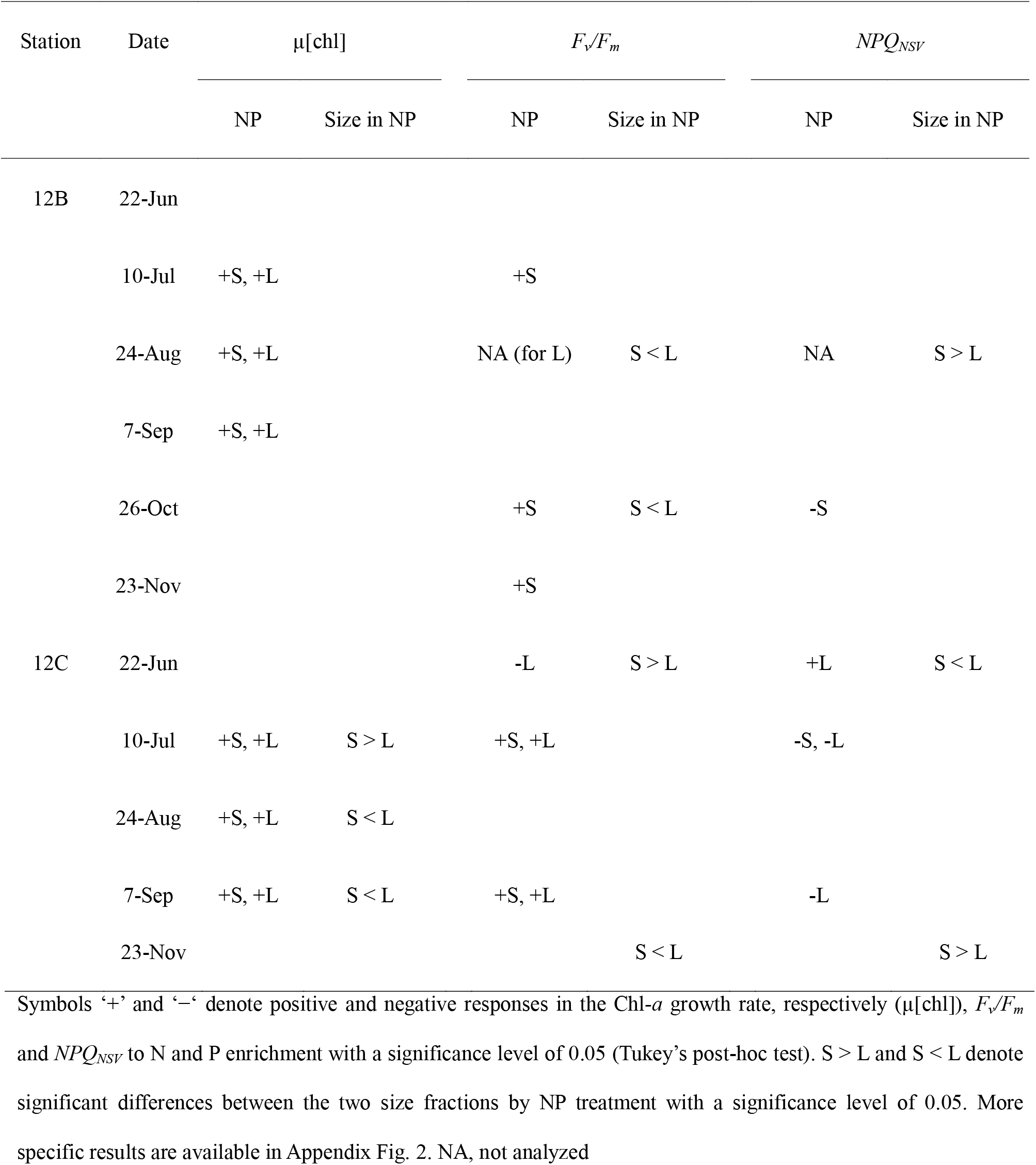
Effects of N and P enrichment (NP) and size fraction on μ[chl], *F_v_/F_m_* and *NPQ_NSV_* of the phytoplankton communities sampled from stations 12B and 12C.

The *F_v_/F_m_* ratio of the small size fraction at station 12B increased with N and P enrichment in July, October and November, while that of the large size fraction did not. In August and October, the *F_v_/F_m_*, ratio of the large size fraction at station 12B remained greater than that of the small fraction even after enrichment. The *F_v_/F_m_* ratio of both size fractions at station 12C increased with enrichment in July and September. In October and November, the *F_v_/F_m_* ratio of the large fractions at station 12C was significantly greater than that of the small fractions in response to N and P enrichment.

The *NPQ_NSV_* value response to N and P enrichment also differed between the size fractions and sampling stations. The *NPQ_NSV_* value of the small size fractions at station 12B decreased with N and P enrichment in October, while there was no change in the large size fraction. However, in August, the *NPQ_NSV_* value of the large size fractions at station 12B was still higher than that of the small size fractions in response to N and P enrichment, but decreased in both size fractions at station 12C in July. In September, the *NPQ_NSV_* value was decreased in only the large size fractions at station 12C with N and P enrichment, while in November, the *NPQ_NSV_* value of the large size fractions at station 12C was lower than that of the small size fractions.

N and P enrichment had only slight effects of on μ[mass] and C_i_, (Appendix Fig. 3). There was also no significant difference in phytoplankton community composition between Con and NP treatments after incubation (PERMANOVA, *p* = 0.38, Appendix Figs. 4, 5).

## Discussion

Over the past few decades, the water temperature and strength of stratification in Lake Biwa have been increasing (Nishino, 2012; Shiga Prefecture, 2018). According to a previous study, the sizes of phytoplankton cells tended to decrease during the stratification period (summer and autumn), but not during the mixing period (winter and spring) (Kishimoto et al., 2013). In this study, the abundance of zygnematophytes, mainly *S. dorsidentiferum* and *M. hardyi*, decreased from July to September during the strong stratification period and increased in October and November in the mixing layer (Fig. 4, Appendix Fig. A1). At the end of the stratification period, the enhanced vertical mixing might increase light fluctuation (Jin et al., 2013) and decrease the sinking loss rate (Zohary et al., 2020). Consequently, the mean cell size of phytoplankton in the large size fractions increased by up to three-fold at station 12B from September to October due to seasonal blooms of large green algae (Fig. 3c).

The results of the present study show for the first time that the sizes of phytoplankton cells in natural lake communities are dependent on *F_v_/F_m_, F_q_′/F_m_′* and *NPQ_NSV_*. The *F_v_/F_m_* and *F_q_′/F_m_′* values of large phytoplankton are greater than those of small phytoplankton in marine communities (Moore et al., 2005; Suggett et al., 2009; Giannini & Ciotti, 2016). Key et al. (2010) demonstrated that the large marine diatom *Coscinodiscus wailesii* has less vulnerability to PSII photoinactivation and, thus, less suppression of *F_v_/F_m_* under excess light than the small marine diatom *Thalassiosira pseudonana*. Previous studies have reported that large phytoplankton cells have a relatively low pigment concentration per cell volume (Agustí, 1991; Álvarez et al., 2016) and, thus, a small absorption cross section per PSII (Morel & Bricaud, 1981; Kirk, 1994; Suggett et al., 2009), which results in lower susceptibility to excess light stress and PSII photodamage. In contrast, small phytoplankton cells are more susceptible to excess light stress and PSII photodamage and, therefore, expend more energy for PSII repair (Key et al., 2010). In this study, the *F_v_/F_m_* values of the large size factions were always greater and the *NPQ_NSV_* values were always lower than those of the small size fractions within the same community (Fig. 5). Moreover, the *F_q_′/F_m_′* ratio of the large size fractions remained greater and the *NPQ_NSV_* value was lower as compared to those of the small size fractions under increased light (Fig. 6, 7). Actually, the *F_v_/F_m_* ratio and *NPQ_NSV_* value were significantly correlated with C_i_, (Fig. 8a, b), but not V_cell_ (Fig. 8c, d). These results indicate that variations in intracellular pigment concentrations may influence susceptibility to light stress rather than cell size (Agusti & Phlips, 1992; Suggett et al., 2009; Campbell & Serôdio, 2020). Indeed, the susceptibility of small phytoplankton to excess light stress decreases with decreasing intracellular pigment concentration in an aggregated colony (Wu et al., 2011).

The advantage of large phytoplankton was pronounced at the end of the stratification period at the pelagic stations in Lake Biwa, as the *F_v_/F_m_* ratio of the small size fractions had improved with N and P enrichment at the end of the stratification period in October and November, but not that of the large size fractions (Table 2). Similarly, the *NPQ_NSV_* values of the small size fractions had decreased in October (Table 2). These results imply that nutrient limitation in PSII photochemistry is size-dependent in the pelagic site in autumn. This phenomenon can be explained by the fluctuation of light and PSII photodamage. At the end of the stratification period, enlargement of the mixing layer causes higher levels of light fluctuation within a water column, which can increase the chance of exposure to excess light for dark-acclimated cells, which induces PSII photodamage (Alderkamp et al., 2010; Helbling et al., 2013). The low-light acclimation reduced the *F_v_/F_m_* ratio and growth rate of algae under fluctuating light conditions due to lower ability to dissipate excess light energy (Yarnold et al., 2016). In autumn, the depth of the mixing layer in Lake Biwa was greater than that of the euphotic zone (Table 2, Fig. 2), thus the phytoplankton became more acclimated to low light or dark conditions. Indeed, the C_i_, of the small size fractions at station 12B increased toward the end of the stratification period (Fig. 3d). These results imply that small phytoplankton tended to acclimate to low light or dark conditions in the enhanced mixing water column at the end of the stratification period and, thus, became sensitive to excess light stress and vulnerable to PSII photoinactivation. Moreover, the rate of PSII repair is influenced by temperature and nutrient availability (Nishiyama & Murata, 2014). Low-temperature stress inhibits D1 protein synthesis, D1 protein degradation in the photodamaged PSII and processing of the D1 preprotein, which generates the mature D1 protein (Nishiyama & Murata, 2014). Synthesis of the D1 protein and xanthophyll, which plays crucial roles in the photoprotective xanthophyll cycle, is also sensitive to nutrient deficiency (Larkum et al., 2003; Shelly et al., 2010). In addition to excess light stress, temperature and nutrient availability might play interactive roles in xanthophyll accumulation or PSII repair (Campbell & Serôdio, 2020). Notably, the μ[chl] of the small size fractions at station 12B did not change in October and November (Table 2), suggesting that small phytoplankton might increase PSII quantum yield without increasing antenna Chl-*a* in order to adapt to excess light. As compared with small phytoplankton, large phytoplankton were relatively less susceptible to PSII photodamage (Suggett et al., 2009; Key et al., 2010) and, thus, conditions were more favorable during the end of the stratification period in Lake Biwa.

The results of the μ[chl] of both small and large phytoplankton suggest that the population growth rates are enhanced by N and P enrichment from July to September (Table 2). Also, the N and P uptake rates of the phytoplankton community at both the beginning and end of the incubation period had increased in August and September (Appendix Fig. 6). However, at station 12C, μ[chl] was greater in the large phytoplankton than the small phytoplankton in response to N and P enrichment in August and September (Table 2). Possibly, cyanobacteria might be favored by N and P enrichment in the large size fractions, which were dominant in these months (Fig. 4d). At the end of the stratification period, N and P limitations to the population growth rate and photochemistry were mitigated, with the exception of small phytoplankton at station 12B (Table 2). In deep lakes and oceans, enhanced vertical mixing increases nutrient availability through transportation from deeper depths to the water surface (Falkowski & Oliver, 2007). Moreover, the autumn typhoon season provides particulate and dissolved nutrients to the lake surface by rainfalls and river discharge, which causes the phytoplankton biomass to rapidly increase (Robarts et al., 1998). An intermittent or fluctuating nutrient supply can favor large phytoplankton (Pinckney et al., 2001; Moore et al., 2008; Bullejos et al., 2010). In marine ecosystems, an intermittent nutrient supply can favor large diatoms due to the presence of relatively large nitrogen storage vacuoles (Litchman et al., 2009). Although many freshwater green algae have storage vacuoles (Tozzi et al., 2004; Becker, 2007; Shebanova et al., 2017), the N and P storage abilities have been confirmed in relatively few species (Shebanova et al., 2017), but not *S. dorsidentiferum* and *M. hardyi* yet. Hence, future studies are warranted to investigate N and P cell storage capacity and interactive effects with light and temperature in these species.

Considering the recent changes in stratification strength, including weakened vertical mixing during the mixing period (Yamada et al., 2021), it can be expected that the mean algal size in Lake Biwa will decrease. However, the speed and the scale of changes in the size structure should also be considered. Indeed, Van de Waal and Litchman (2020) pointed out that the increasing light availability due to shrinking of the mixing layer can benefit large diatoms. In regard to river phytoplankton, increases in the frequency of sudden flood events will result in increased spring blooms of large phytoplankton (Abonyi et al., 2018). The grazing effects of crustacean zooplankton likely also play important roles (Lampert et al., 1986; Sunda & Hardison, 2010; Branco et al., 2020). Yvon-Durocher et al. (2015) revealed that enhanced zooplankton grazing due to global warming has increased the mean cell size of phytoplankton communities. Future studies should focus on whether light environments, mixing events and predation effects can mitigate the shrinking of phytoplankton cell size with the warming of lake waters. Because phytoplankton size structure is an important factor in the material cycle (Ray et al., 2001; Law et al., 2009) and trophic structure (Kazama et al., 2021b) in lakes, it is necessary to determine the response of local phytoplankton communities to global climate change in order to develop better conservation and management plans for aquatic ecosystems in the future.

## Acknowledgements

We would like to thank Hirokazu Teraishi for assisting with the chemical analysis. This study was financially supported by the Collaborative Research Fund from Shiga Prefecture entitled ‘Study on water quality and lake-bottom environment for protection of the soundness of water environment’ under the Japanese Grant for Regional Revitalization and the Environment Research and Technology Development Fund (grant no. 5-1607) of the Ministry of the Environment, Japan (https://www.kantei.go.jp/jp/singi/tiiki/tiikisaisei/souseikoufukin.html).

## Declarations

### Availability of data and material

All data generated or analyzed during this study are included in this published article.

### Authors’ contributions

Conceptualization (TK, KH), Methodology (TK, KH, KK), Resources (KH, TN), Investigation (TK, KH, TN, KS), Formal analysis (TK), Writing-original draft (TK), Writing-review & editing (TK, KH, TN, KS, AI, KK), Supervision (AI).

### Ethics approval

Not applicable

### Consent to participate

Not applicable

### Consent for publication

Not applicable

## Supporting information

**Appendix Fig. 1-6**

## Abbreviations

*F′*: Fluorescence yield under actinic light
*F_m_*: Maximum PSII fluorescence yield in dark-adapted state
*F_m_′*: Maximum PSII fluorescence yield in light-adapted state
*F_O_*: Minimum PSII fluorescence yield in dark-adapted state
*F_O_′*: Minimum PSII fluorescence yield in light-adapted state
*F_q_*: Variable PSII fluorescence yield in light-adapted state (*F_m_ - F*)
*F_v_*: Maximum variable PSII fluorescence yield in dark-adapted state (*F_m_ - F_O_*)
*F_v_′*: Variable PSII fluorescence yield under actinic light (*F_m_′ - F_O_′*)
*F_v_/F_m_*: Maximum PSII photochemical efficiency in dark-adapted state
*F_q_′/F_m_′*: Effective PSII photochemical efficiency in light-adapted state
*NPQ_NSV_*: Normalized Stern-Volmer coefficient of quenching (*F_O_′/F_v_′*)

