## Appendix_Fig for "Size-dependent susceptibility of lake phytoplankton to light stress: An implication for succession of large green algae in a deep oligotrophic lake"

### Slide 1
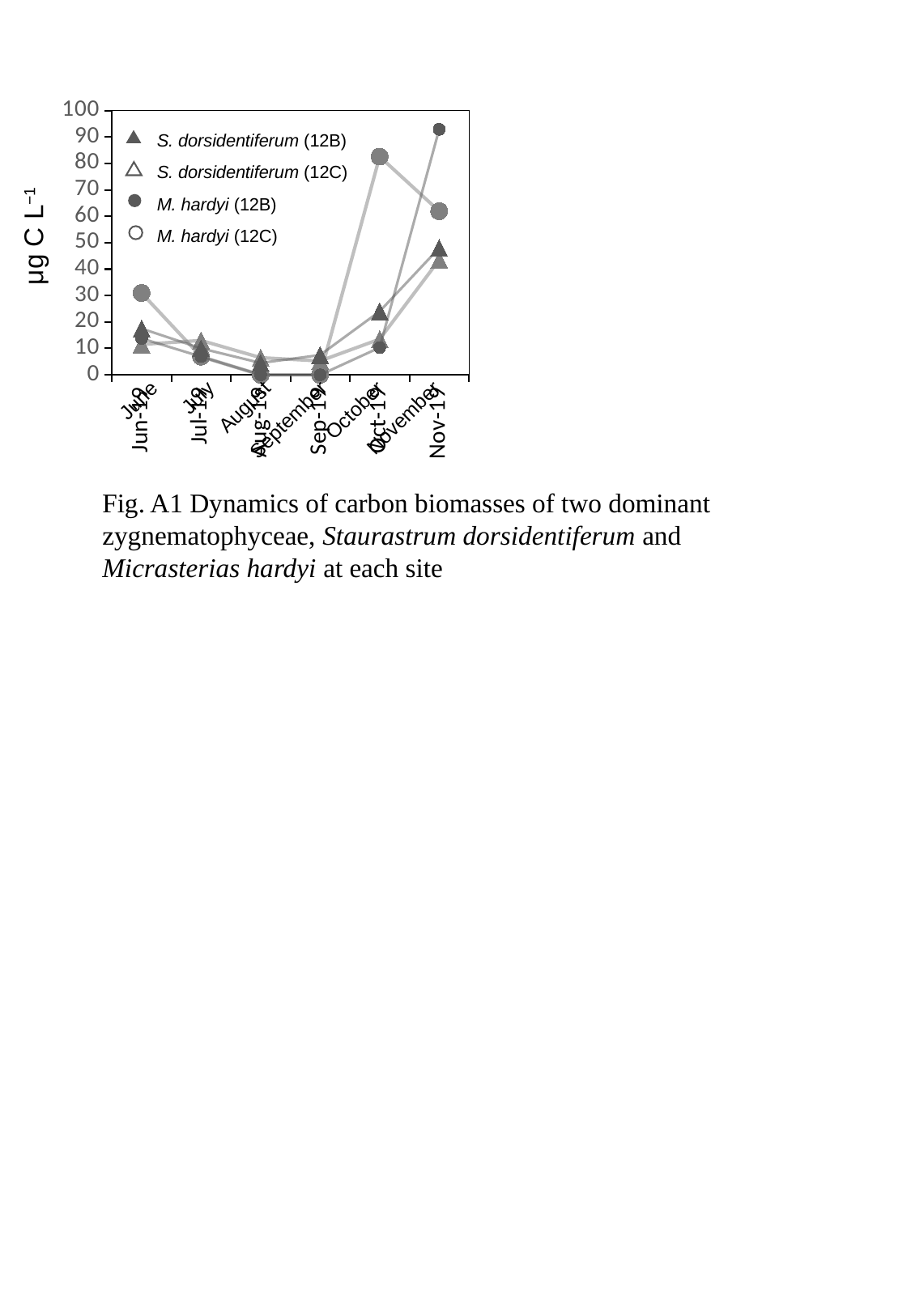

#### Chart
| Category | | | | |
|---|---|---|---|---|
| | 11.503882647228885 | 30.96511 | 17.505908376217867 | 13.762271111111112 |
| | 13.00438907947613 | 6.881135555555556 | 10.003376214981637 | 6.881135555555556 |
| | 6.502194539738065 | 0.0 | 4.501519296741737 | 0.0 |
| | 5.25177251286536 | 0.0 | 7.502532161236228 | 0.0 |
| | 13.504557890225211 | 82.57362666666667 | 24.008102915955934 | 10.321703333333334 |
| | 43.51468653517013 | 61.93022 | 48.01620583191187 | 92.89533 |S. dorsidentiferum (12B)
S. dorsidentiferum (12C)
M. hardyi (12B)
M. hardyi (12C)
μg C L^−1^
June
July
August
September
October
November
Fig. A1 Dynamics of carbon biomasses of two dominant zygnematophyceae, Staurastrum dorsidentiferum and Micrasterias hardyi at each site

### Slide 2
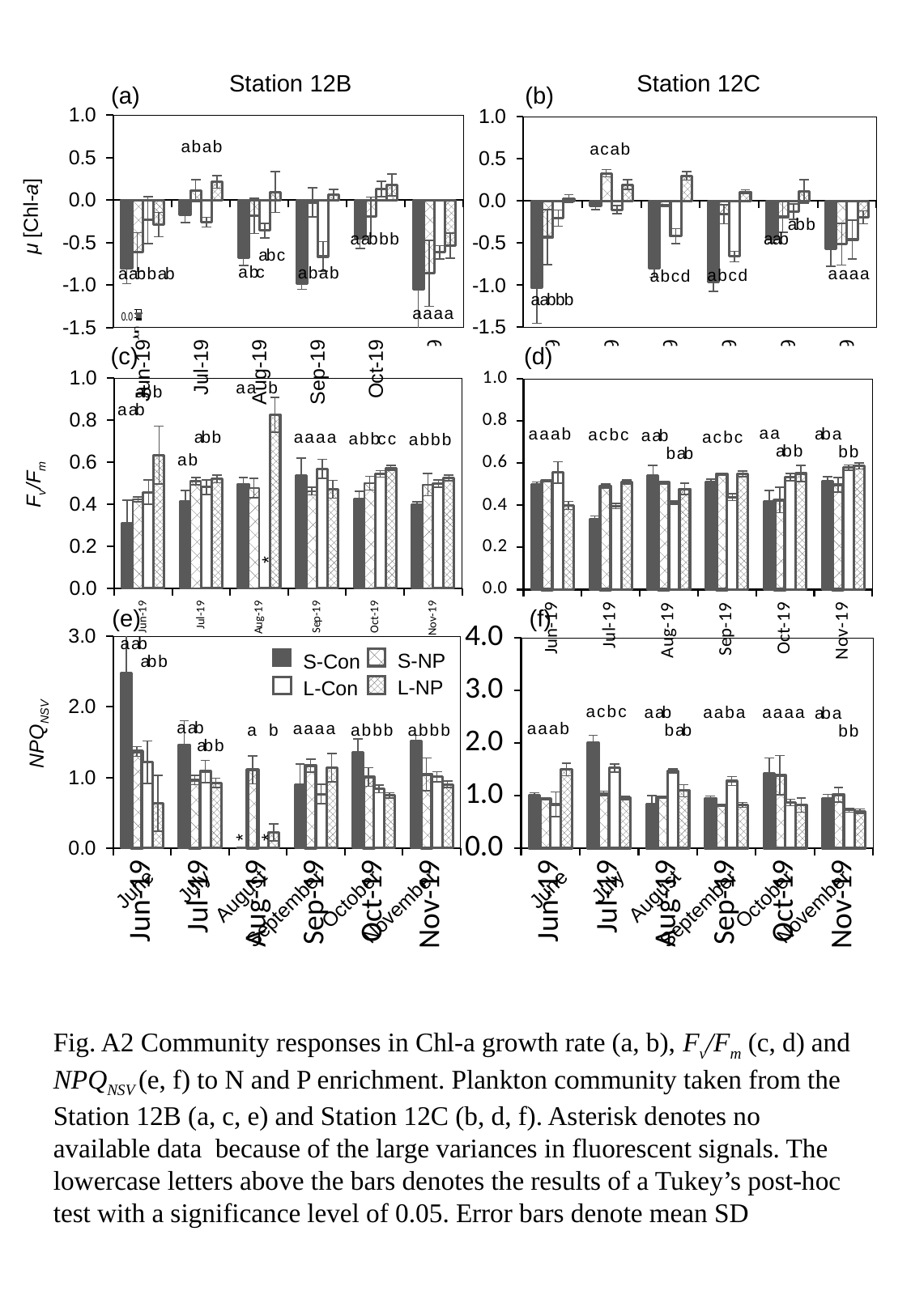

#### Chart
| Category | 12C | 12C | 12C | 12C |
|---|---|---|---|---|
| 43638 | -1.0255664318407893 | -0.42990649498949957 | -0.20413566567546834 | 0.030520134919160335 |
| 43656 | -0.06207376711340843 | 0.33005123916288726 | -0.10365561241947858 | 0.19455939239682518 |
| 43701 | -0.795939438893931 | -0.05387144141212883 | -0.41778674200792815 | 0.29917644137716504 |
| 43715 | -0.9595913396723805 | -0.15698988928346488 | -0.6591971322590465 | 0.10844262687297528 |
| 43764 | -0.44843328187970194 | -0.18876665411144342 | -0.1251936939171329 | 0.11393841906253421 |
| 43792 | -0.5706338324889026 | -0.5130846908078256 | -0.4580171732574372 | -0.19356773131721153 |
#### Chart
| Category | 12B | 12B | 12B | 12B |
|---|---|---|---|---|
| 43638 | -0.8022775865320657 | -0.6136430951375321 | -0.23603060896279204 | -0.2898685227969942 |
| 43656 | -0.17024841100391905 | 0.11126643446813474 | -0.2612597667551711 | 0.21491882738844081 |
| 43701 | -0.6796156625624455 | -0.18675932259316355 | -0.3595881092506228 | 0.094708173928354 |
| 43715 | -0.9857263431248935 | -0.027830243353151162 | -0.6655382953529837 | 0.06501622446805637 |
| 43764 | -0.4519334986164807 | -0.1933843224546116 | 0.13169295838453368 | 0.1777477466305523 |
| 43792 | -1.0511366521456826 | -0.8630426729952186 | -0.615924159304475 | -0.5371118598217438 |Station 12C
Station 12B
(a)
(b)
a b a b
a c a b
μ [Chl-a]
ab b
a ab b b
a ab
ab c
 a bc
a b a b
a ab b ab
a a a a
a b c d
a b c d
a ab b b
a a a a
#### Chart
| Category | 12C | 12C | 12C | 12C |
|---|---|---|---|---|
| 43638 | 0.49899999999999994 | 0.5166666666666667 | 0.5559999999999999 | 0.39999999999999997 |
| 43656 | 0.3333333333333333 | 0.49099999999999994 | 0.3960000000000001 | 0.5116666666666667 |
| 43701 | 0.5436666666666666 | 0.508 | 0.4126666666666667 | 0.47833333333333333 |
| 43715 | 0.5123333333333333 | 0.549 | 0.43933333333333335 | 0.5489999999999999 |
| 43764 | 0.4176666666666667 | 0.425 | 0.535 | 0.5513333333333333 |
| 43792 | 0.5146666666666667 | 0.49733333333333335 | 0.5796666666666667 | 0.5869999999999999 |
#### Chart
| Category | 12B | 12B | 12B | 12B |
|---|---|---|---|---|
| 43638 | 0.3075 | 0.4226666666666667 | 0.4566666666666667 | 0.6343333333333333 |
| 43656 | 0.41133333333333333 | 0.5093333333333333 | 0.48100000000000004 | 0.5203333333333333 |
| 43701 | 0.495 | 0.4776666666666667 | 0.0 | 0.8256666666666667 |
| 43715 | 0.5363333333333333 | 0.4623333333333333 | 0.5690000000000001 | 0.4706666666666666 |
| 43764 | 0.426 | 0.4996666666666667 | 0.5443333333333333 | 0.573 |
| 43792 | 0.39766666666666667 | 0.49333333333333335 | 0.498 | 0.5256666666666667 |(c)
(d)
#### Chart
| Category | 12C | 12C | 12C | 12C |
|---|---|---|---|---|
| 43638 | 0.49899999999999994 | 0.5166666666666667 | 0.5559999999999999 | 0.39999999999999997 |
| 43656 | 0.3333333333333333 | 0.49099999999999994 | 0.3960000000000001 | 0.5116666666666667 |
| 43701 | 0.5436666666666666 | 0.508 | 0.4126666666666667 | 0.47833333333333333 |
| 43715 | 0.5123333333333333 | 0.549 | 0.43933333333333335 | 0.5489999999999999 |
| 43764 | 0.4176666666666667 | 0.425 | 0.535 | 0.5513333333333333 |
| 43792 | 0.5146666666666667 | 0.49733333333333335 | 0.5796666666666667 | 0.5869999999999999 |a a b
a ab
a a
a a a b
ab a
a c b c
a ab
a a a a
a c b c
a b bc c
a b b b
ab b
b b
a b
F_v_/F_m_
*
(e)
(f)
#### Chart
| Category | 12C | 12C | 12C | 12C |
|---|---|---|---|---|
| 43638 | 1.0043346693386772 | 0.9343333333333333 | 0.8320000000000001 | 1.5008132208975475 |
| 43656 | 2.003 | 1.038 | 1.527 | 0.9540000000000001 |
| 43701 | 0.843 | 0.9683333333333333 | 1.4683333333333335 | 1.095 |
| 43715 | 0.9533333333333335 | 0.8220000000000001 | 1.2793333333333334 | 0.822 |
| 43764 | 1.4193333333333333 | 1.3870000000000002 | 0.8700000000000001 | 0.82 |
| 43792 | 0.9456666666666665 | 1.0166666666666666 | 0.7253333333333334 | 0.7040000000000001 |
#### Chart
| Category | 12B | 12B | 12B | 12B |
|---|---|---|---|---|
| 43638 | 2.4745 | 1.3673333333333335 | 1.2143333333333333 | 0.6333333333333333 |
| 43656 | 1.4606666666666666 | 0.9653333333333333 | 1.086 | 0.9226666666666666 |
| 43701 | 0.0 | 1.1066666666666667 | 0.0 | 0.21966666666666665 |
| 43715 | 0.8943333333333333 | 1.1663333333333332 | 0.7643333333333334 | 1.1376666666666668 |
| 43764 | 1.3556666666666668 | 1.0073333333333332 | 0.8383333333333333 | 0.7456666666666667 |
| 43792 | 1.517666666666667 | 1.0436666666666667 | 1.0090000000000001 | 0.902 |
a ab
S-NP
L-NP
S-Con
L-Con
ab b
a c b c
a ab
a a b a
a a a a
ab a
a ab
a a a a
a a a b
 a b
a b b b
b ab
a b b b
b b
NPQ_NSV_
ab b
*
*
June
July
August
September
October
November
June
July
August
September
October
November
Fig. A2 Community responses in Chl-a growth rate (a, b), F_v_/F_m_ (c, d) and NPQ_NSV_ (e, f) to N and P enrichment. Plankton community taken from the Station 12B (a, c, e) and Station 12C (b, d, f). Asterisk denotes no available data because of the large variances in fluorescent signals. The lowercase letters above the bars denotes the results of a Tukey’s post-hoc test with a significance level of 0.05. Error bars denote mean SD

### Slide 3
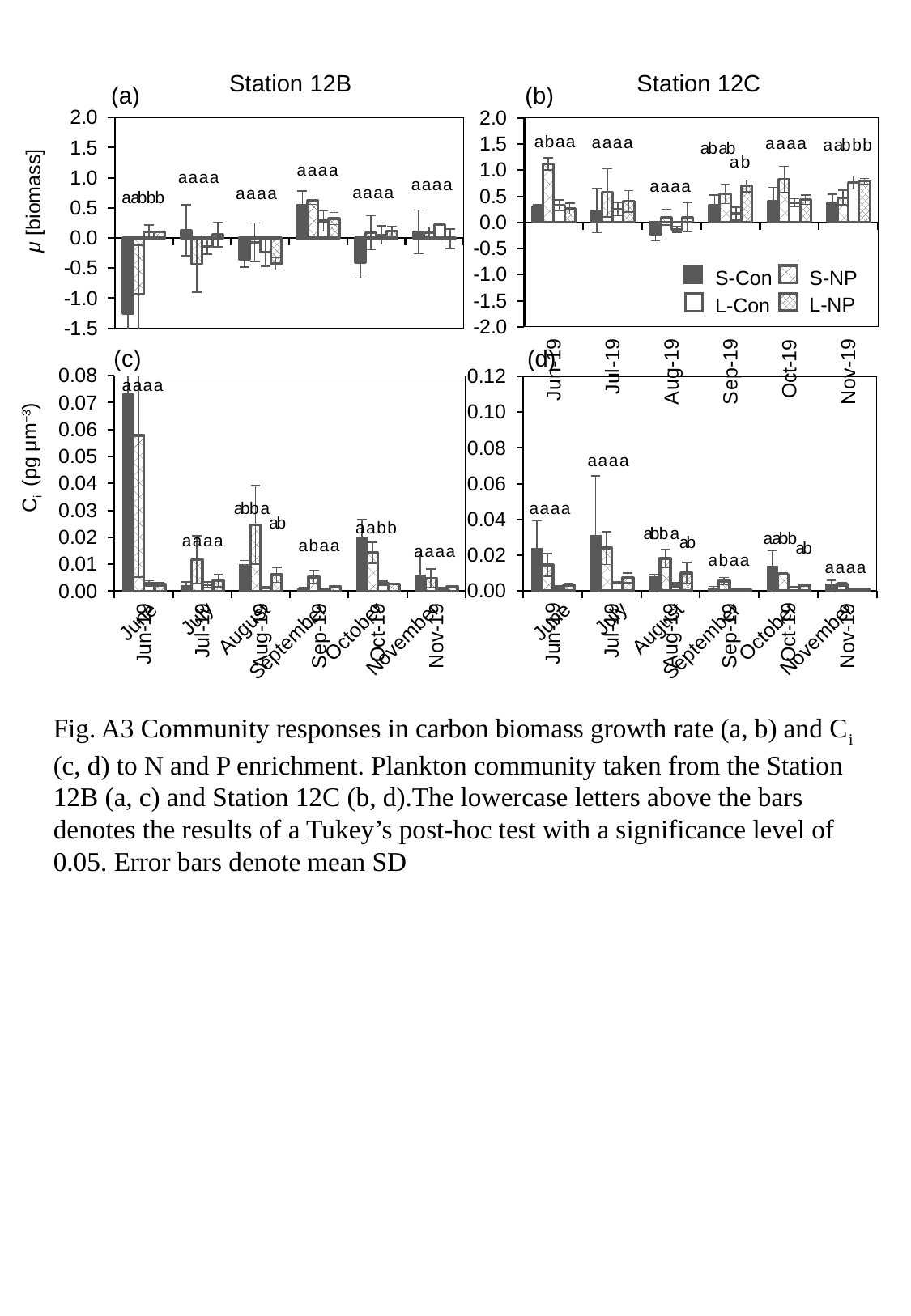

Station 12C
Station 12B
(a)
(b)
#### Chart
| Category | 12B | 12B | 12B | 12B |
|---|---|---|---|---|
| 43638 | -1.251139844805767 | -0.9326321762929742 | 0.10213197451252247 | 0.0990992627480383 |
| 43656 | 0.1302316300074464 | -0.4377411157963496 | -0.14489266466220807 | 0.05654370043814035 |
| 43701 | -0.3496543981506079 | -0.06951803119834427 | -0.2390815140040056 | -0.4293808593451229 |
| 43715 | 0.5374210618390705 | 0.6189065974015989 | 0.2822875806433574 | 0.32220775859786716 |
| 43764 | -0.4056979558973441 | 0.0873204449979361 | 0.05104540833180562 | 0.10990068695162886 |
| 43792 | 0.10184749126443195 | 0.08551895233253033 | 0.22279526517059642 | -0.013992568076074232 |
#### Chart
| Category | 12C | 12C | 12C | 12C |
|---|---|---|---|---|
| 43638 | 0.3052378760083337 | 1.1190185452205197 | 0.3333159520926381 | 0.26336432698771817 |
| 43656 | 0.227201757385426 | 0.5729132582991604 | 0.2525699305491298 | 0.4020425751436822 |
| 43701 | -0.22771703230214788 | 0.10302242277154876 | -0.13232077030183986 | 0.1018194252075475 |
| 43715 | 0.32692055929591135 | 0.5492151667720279 | 0.16662486060949713 | 0.7000764701867505 |
| 43764 | 0.4031603138309013 | 0.8243151643482056 | 0.37860954802041663 | 0.4327889063881352 |
| 43792 | 0.3758458109204201 | 0.4739187547894738 | 0.7654708590879123 | 0.7888125943920873 |a b a a
a a a a
a a a a
a ab b b
ab ab
 a b
a a a a
a a a a
a a a a
a a a a
a a a a
a a a a
a ab b b
μ [biomass]
S-NP
L-NP
S-Con
L-Con
(c)
(d)
#### Chart
| Category | 12B | 12B | 12B | 12B |
|---|---|---|---|---|
| 43638 | 0.07322569374245368 | 0.05777099603692618 | 0.0028859594086083577 | 0.0025700120468744795 |
| 43656 | 0.0020236487532935515 | 0.01173330881229538 | 0.0023708721335320548 | 0.003921932913709024 |
| 43701 | 0.009819090989226639 | 0.02464563646826685 | 0.0013303967141447166 | 0.006038676063932404 |
| 43715 | 0.0008686284247621043 | 0.00531759453761414 | 0.00045200675968853373 | 0.001659793179513469 |
| 43764 | 0.02025266421590789 | 0.014247925009323514 | 0.002882944479475933 | 0.0026312335318452364 |
| 43792 | 0.006062083844697037 | 0.004814955561653091 | 0.0009455277189776569 | 0.0016589468921605268 |
#### Chart
| Category | 12C | 12C | 12C | 12C |
|---|---|---|---|---|
| 43638 | 0.02377069608180957 | 0.014594386543913283 | 0.0020059420378002167 | 0.003405433671935768 |
| 43656 | 0.031191138817220687 | 0.023920866416068248 | 0.004693812668854244 | 0.007341391850733803 |
| 43701 | 0.008068320188832563 | 0.01807473210127516 | 0.0034662475304146185 | 0.009981934752913578 |
| 43715 | 0.0015853041752628267 | 0.005583475027956478 | 0.00045661614698864296 | 0.0007548725460243925 |
| 43764 | 0.013760719628377521 | 0.00936682178855884 | 0.00202887636113791 | 0.003109909699230842 |
| 43792 | 0.00386146894811127 | 0.0037471013070618166 | 0.000887594481133592 | 0.001205889253990929 |a a a a
C_i_ (pg μm^−3^)
a a a a
a a a a
ab b a
ab
a a b b
ab b a
a ab b
a a a a
a b a a
a a a a
a b a a
a a a a
June
July
August
September
October
November
June
July
August
September
October
November
Fig. A3 Community responses in carbon biomass growth rate (a, b) and C_i_ (c, d) to N and P enrichment. Plankton community taken from the Station 12B (a, c) and Station 12C (b, d).The lowercase letters above the bars denotes the results of a Tukey’s post-hoc test with a significance level of 0.05. Error bars denote mean SD

### Slide 4
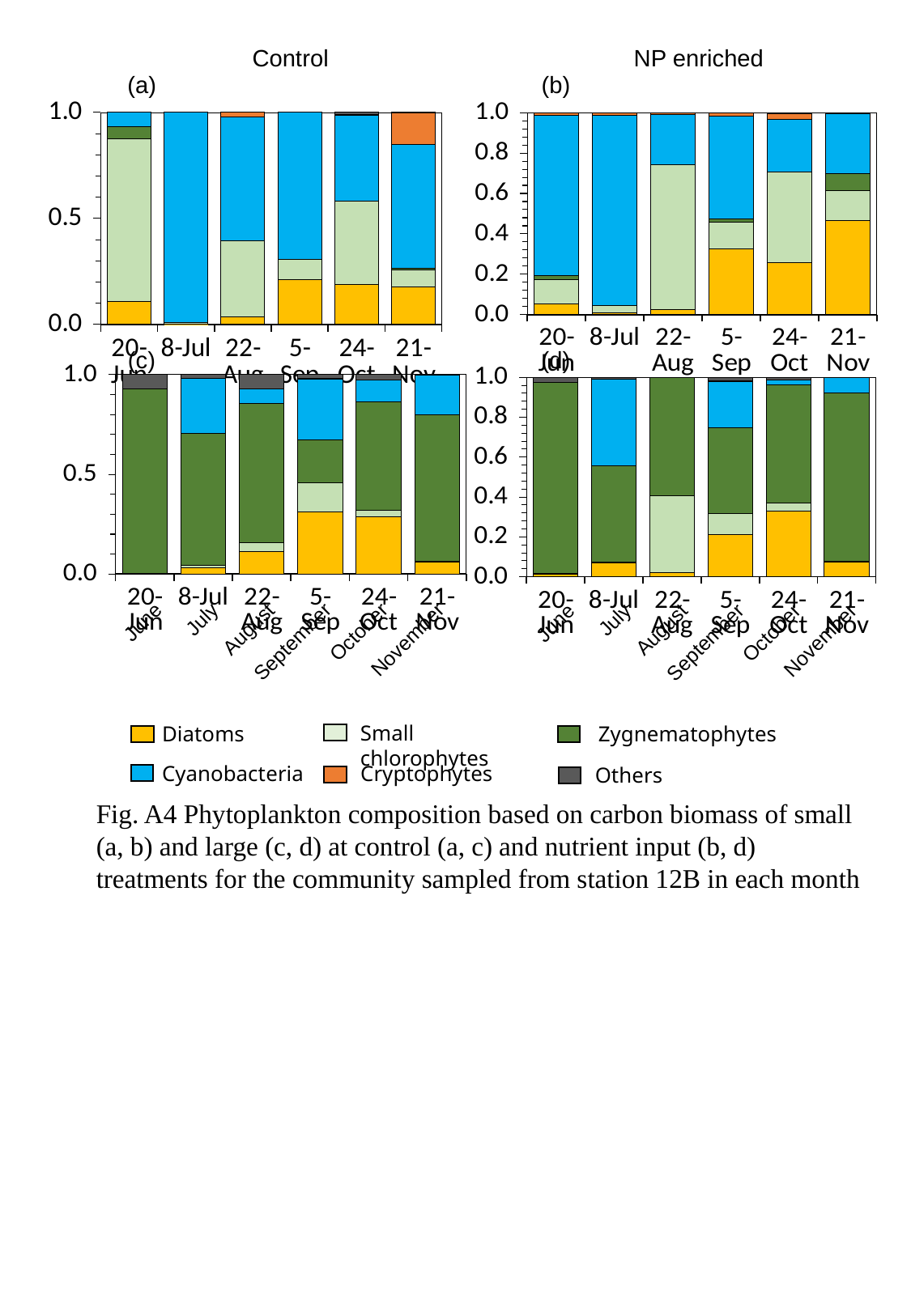

NP enriched
Control
(a)
(b)
#### Chart
| Category | Diatoms | Chlorophytes | Zygnematophytes | Cyanophytes | Cryptophytes | others |
|---|---|---|---|---|---|---|
| 43636 | 0.09770256738336346 | 0.7015290480982795 | 0.05292012678176985 | 0.06161025015057131 | 0.0 | 0.0 |
| 43654 | 0.08623406226477537 | 0.4943670773841748 | 0.0 | 71.95744687362078 | 0.0 | 0.0 |
| 43699 | 0.1430360110206161 | 1.4111235336581371 | 0.0 | 2.2921872262001 | 0.07776244584043458 | 0.0 |
| 43713 | 5.366503467868786 | 2.3764984454230973 | 0.0 | 17.621923513098903 | 0.0 | 0.0 |
| 43762 | 1.1661870608366798 | 2.3894046367064203 | 0.0 | 2.488810991822609 | 0.017053238747495078 | 0.06610720783088023 |
| 43790 | 1.2595180214907813 | 0.5566742229233415 | 0.06614446342387865 | 4.149095847835352 | 1.0700991882843522 | 0.009476420215553996 |
#### Chart
| Category | Diatoms | Chlorophytes | Zygnematophytes | Cyanophytes | Cryptophytes | others |
|---|---|---|---|---|---|---|
| 43636 | 0.1421910876776724 | 0.3099849743495835 | 0.05292012678176985 | 2.082286314705662 | 0.03888122292021729 | 0.0 |
| 43654 | 0.2347693288155959 | 0.8735177046162431 | 0.0 | 22.661477891515506 | 0.3117107225464322 | 0.0 |
| 43699 | 0.18194596495288093 | 5.440637378104256 | 0.0 | 1.8599269970682322 | 0.07776244584043458 | 0.0 |
| 43713 | 9.04466832090987 | 3.733214279116659 | 0.5001688107490819 | 14.154485701384205 | 0.506139157481225 | 0.0 |
| 43762 | 4.316330822026237 | 7.547657020260982 | 0.0 | 4.393476559669106 | 0.5070808437314559 | 0.055447766020657034 |
| 43790 | 2.70431907490203 | 0.8501256524586328 | 0.48281722236781704 | 1.7255831436906106 | 0.008526619373747539 | 0.014214630323330993 |(c)
(d)
#### Chart
| Category | Diatoms | Chlorophytes | Zygnematophytes | Cyanophytes | Cryptophytes | others |
|---|---|---|---|---|---|---|
| 43636 | 0.23199441384701605 | 0.039905533556411736 | 57.56739662735462 | 0.0 | 0.0 | 4.337459527935364 |
| 43654 | 1.3423477233667487 | 0.5225844634228624 | 28.339969786423378 | 11.957020649592806 | 0.0 | 0.7229099213225606 |
| 43699 | 1.1598319190688535 | 0.4347976858607831 | 7.0453155539681305 | 0.7543350021111602 | 0.0 | 0.7229099213225606 |
| 43713 | 11.554023386799612 | 5.4165488590386905 | 7.9749264397913615 | 11.435421495062327 | 0.07776244584043458 | 0.7229099213225606 |
| 43762 | 36.86651283206944 | 4.235795828919501 | 70.51496904894252 | 14.007875137988423 | 0.0 | 3.386256450381752 |
| 43790 | 11.660021738833157 | 0.18358524681967817 | 139.85731591999885 | 38.18976452640375 | 0.12616169691144016 | 0.0 |
#### Chart
| Category | Diatoms | Chlorophytes | Zygnematophytes | Cyanophytes | Cryptophytes | others |
|---|---|---|---|---|---|---|
| 43636 | 0.6693346361006443 | 0.4361012879121127 | 58.66503756903595 | 0.0 | 0.03888122292021729 | 1.4458198426451212 |
| 43654 | 4.687038595929034 | 0.30346803537946543 | 31.993485213131198 | 28.841355386291724 | 0.0 | 0.7229099213225606 |
| 43699 | 0.13169302555254817 | 2.5464212117933247 | 3.9078867012901757 | 0.0 | 0.0 | 0.0 |
| 43713 | 8.278401478140387 | 4.148065914297952 | 16.95248846415977 | 9.117979236984967 | 0.04547530332665354 | 0.7229099213225606 |
| 43762 | 46.9437323038134 | 5.878926309561184 | 84.5718163666582 | 3.2483329318320906 | 0.011368825831663383 | 1.9602641451033103 |
| 43790 | 9.008595153491276 | 0.6177481561232132 | 103.5307544913306 | 9.514552577151038 | 0.025579858121242614 | 0.0 |June
July
August
September
October
November
June
July
August
September
October
November
Small chlorophytes
Diatoms
Zygnematophytes
Cyanobacteria
Cryptophytes
Others
Fig. A4 Phytoplankton composition based on carbon biomass of small (a, b) and large (c, d) at control (a, c) and nutrient input (b, d) treatments for the community sampled from station 12B in each month

### Slide 5
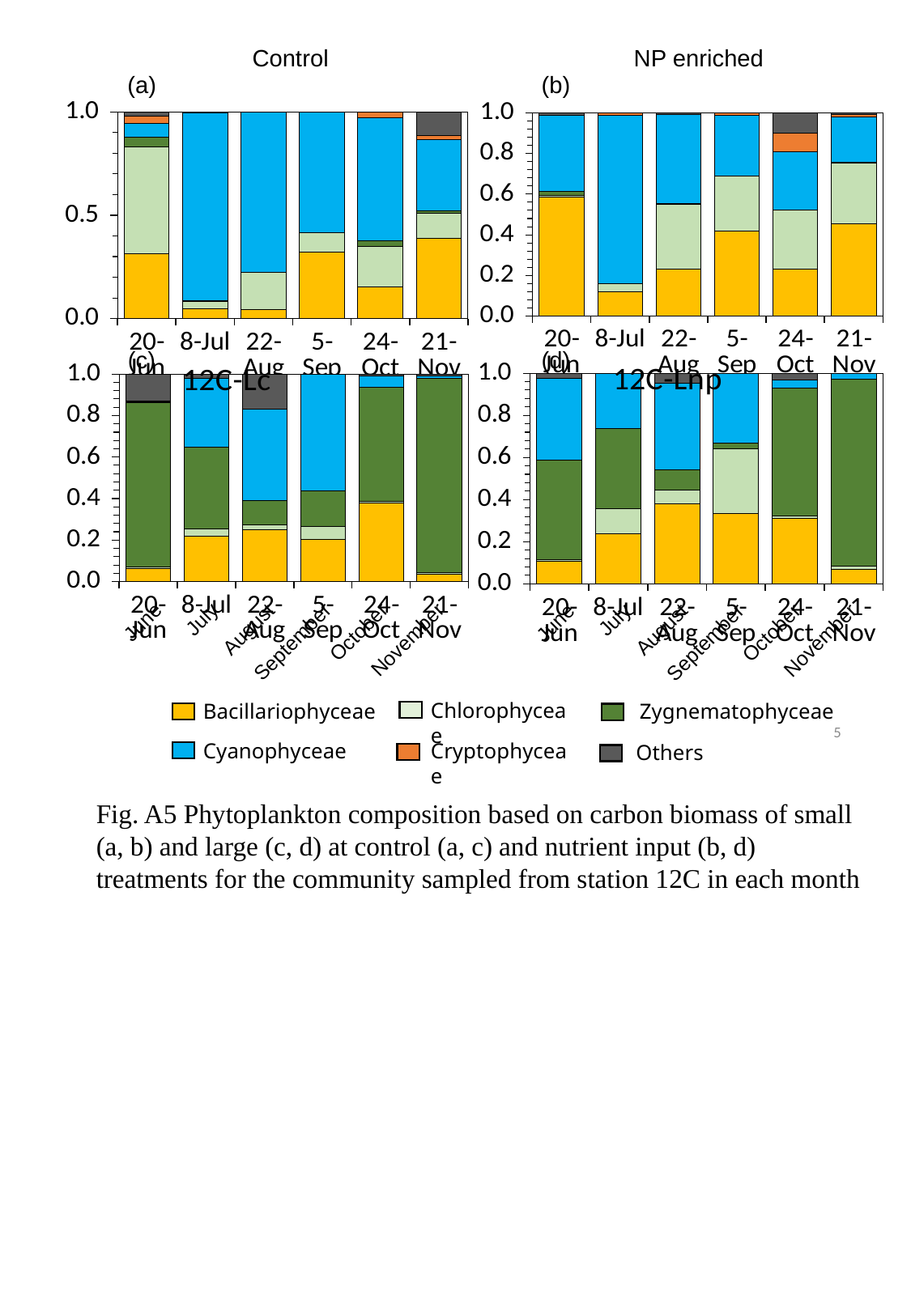

NP enriched
Control
(a)
(b)
#### Chart
| Category | Diatoms | Chlorophytes | Zygnematophytes | Cyanophytes | Cryptophytes | others |
|---|---|---|---|---|---|---|
| 43636 | 10.491668549154939 | 0.11546130886001538 | 0.3175207606906191 | 6.745075721514939 | 0.0 | 0.19832162349264068 |
| 43654 | 8.416093578707189 | 2.743880742496419 | 0.0 | 58.360479743603086 | 0.9344712284546026 | 0.0 |
| 43699 | 3.6357342051760804 | 4.962961469571816 | 0.09941712225707759 | 6.826060244058922 | 0.0 | 0.159973633630944 |
| 43713 | 30.665617919953515 | 19.565994922287448 | 0.0 | 21.892648732128404 | 0.7564797337500239 | 0.12174162459594727 |
| 43762 | 5.6421541911832165 | 7.040220060495289 | 0.0 | 6.966918750923427 | 2.2251523854457527 | 2.4866552555420487 |
| 43790 | 14.939447865901371 | 9.81142491257488 | 0.1322889268477573 | 7.34791131147855 | 0.38874930380284095 | 0.2345414003349614 |
#### Chart
| Category | Diatoms | Chlorophytes | Zygnematophytes | Cyanophytes | Cryptophytes | others |
|---|---|---|---|---|---|---|
| 43636 | 2.8990357994975433 | 4.822976878169498 | 0.4233610142541588 | 0.6455488404743579 | 0.312371661731126 | 0.18457429322159033 |
| 43654 | 2.4322603034341026 | 1.6677583576191406 | 0.18522044373619448 | 45.081183876306056 | 0.15552489168086917 | 0.0 |
| 43699 | 0.33756510414520924 | 1.460533938463179 | 0.0 | 6.233639399014336 | 0.0 | 0.0 |
| 43713 | 15.141166440662872 | 4.482568295285961 | 0.0 | 27.536689340014988 | 0.022737651663326766 | 0.0 |
| 43762 | 1.659420324594137 | 2.101482499535441 | 0.28438383209618107 | 6.384874111007552 | 0.3106452282032063 | 0.0 |
| 43790 | 10.620551480643167 | 3.3090732672253345 | 0.3254719271550029 | 9.329781871051388 | 0.6240194588782262 | 3.0510182360845977 |(c)
(d)
#### Chart: 12C-Lnp
| Category | Diatoms | Chlorophytes | Zygnematophytes | Cyanophytes | Cryptophytes | others |
|---|---|---|---|---|---|---|
| 43636 | 12.202990379458328 | 1.2717772880782106 | 54.65417777296426 | 45.09863171836961 | 0.0 | 2.8916396852902424 |
| 43654 | 31.525025490033432 | 15.966182329423889 | 50.984501977132034 | 34.943564155634085 | 0.0 | 0.0 |
| 43699 | 17.373394126432334 | 3.077121318068023 | 4.408055512039258 | 18.802028696891238 | 0.0 | 2.168729763967682 |
| 43713 | 124.74338853397721 | 115.27964157908957 | 9.474819424940941 | 123.89107778041341 | 0.23328733752130373 | 0.12174162459594727 |
| 43762 | 57.584678951979335 | 2.5858482595644285 | 112.5813077406962 | 7.81043864940679 | 0.03888122292021729 | 5.617248596982916 |
| 43790 | 10.990717582760373 | 2.6439008633281325 | 144.06883084105095 | 4.357447761566829 | 0.08390169250156855 | 0.12174162459594727 |
#### Chart: 12C-Lc
| Category | Diatoms | Chlorophytes | Zygnematophytes | Cyanophytes | Cryptophytes | others |
|---|---|---|---|---|---|---|
| 43636 | 5.066506955339273 | 0.5991090455865465 | 65.79354642154458 | 0.3248666024503725 | 0.07842338502512841 | 10.884229361370389 |
| 43654 | 21.217130287219216 | 3.4229364864147294 | 38.338162435758505 | 32.318290295433656 | 0.0 | 2.168729763967682 |
| 43699 | 6.477736295988092 | 0.5656505036621903 | 3.0010128644944913 | 11.391099090132931 | 0.0 | 4.337459527935364 |
| 43713 | 26.244080676228094 | 7.9666514805363855 | 22.443755613298766 | 72.55608727584877 | 0.0 | 0.0 |
| 43762 | 62.892397394870954 | 1.8847060702959493 | 91.36405828929361 | 9.166414824941228 | 0.0 | 1.5675614672410685 |
| 43790 | 5.399414133959745 | 1.3215544047436671 | 147.3605296372676 | 2.1750631490472814 | 0.0 | 1.2028308222153925 |June
July
August
September
October
November
June
July
August
September
October
November
Chlorophyceae
Bacillariophyceae
Zygnematophyceae
Cyanophyceae
Cryptophyceae
Others
5
Fig. A5 Phytoplankton composition based on carbon biomass of small (a, b) and large (c, d) at control (a, c) and nutrient input (b, d) treatments for the community sampled from station 12C in each month

### Slide 6
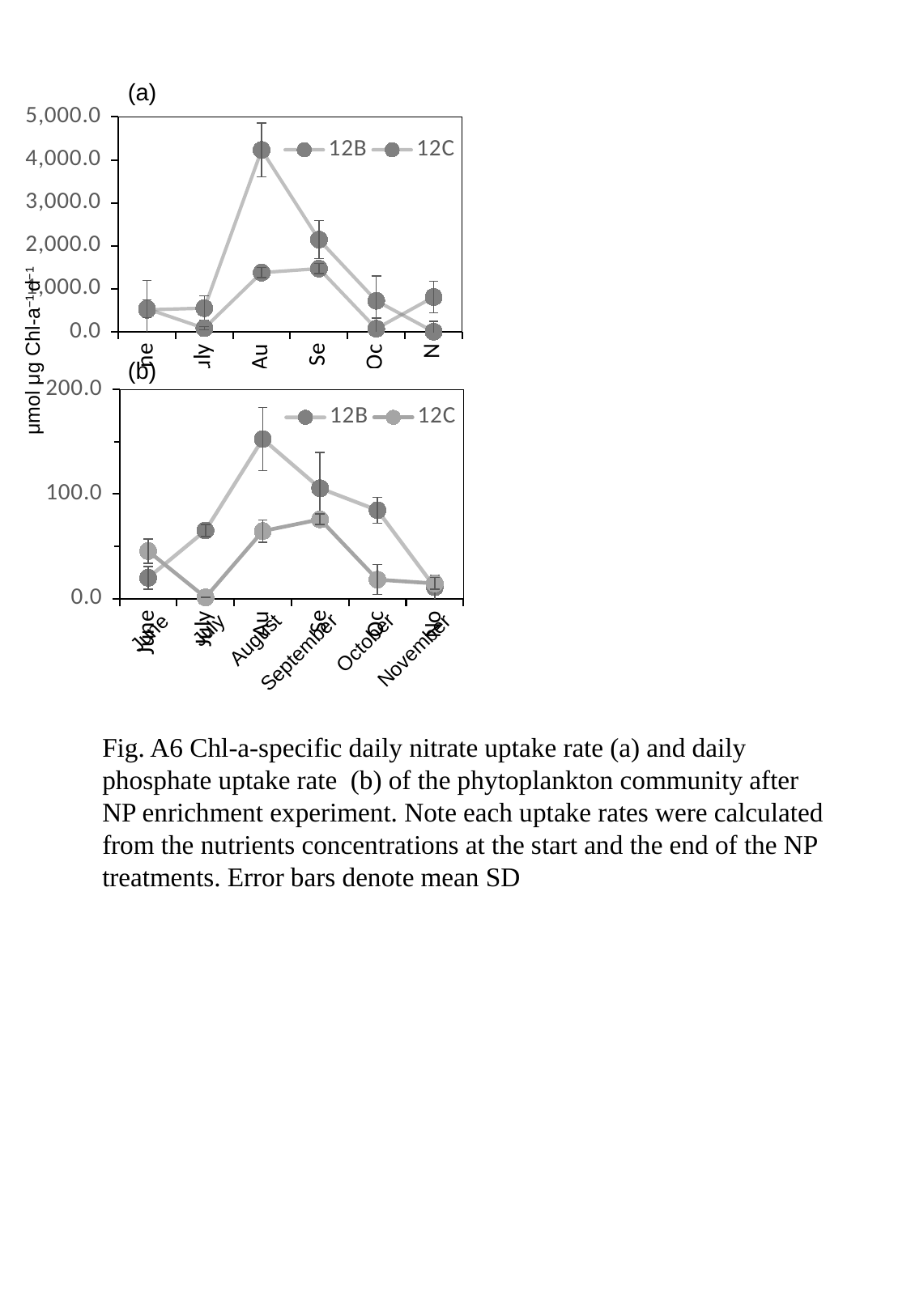

#### Chart
| Category | | |
|---|---|---|
| June | 25.66359967943204 | 11.745612266643198 |
| July | 8.422239089396761 | 71.74362026018227 |
| August | 27.708129292010877 | 21.30469458847278 |
| September | 20.315798759981508 | 19.392597802006645 |
| October | 8.513245070585198 | 3.900522166212639 |
| November | 0.1 | 55.68868109686082 |(a)
#### Chart
| Category | | |
|---|---|---|
| June | 509.9771353439956 | 534.1415028113854 |
| July | 548.8301278101254 | 81.05967439037478 |
| August | 4229.941634625833 | 1375.6149468215178 |
| September | 2143.6936618021487 | 1469.425070988842 |
| October | 719.9215556363716 | 70.83454424744035 |
| November | 0.3965069327795197 | 807.4713366287561 |June
July
August
September
October
November
μmol μg Chl-a^−1^ d^−1^
(b)
#### Chart
| Category | | |
|---|---|---|
| June | 19.871613558277023 | 45.47583307583835 |
| July | 65.16439654403531 | 1.1298520216349177 |
| August | 152.66067189333702 | 64.56863021945476 |
| September | 105.51855170099647 | 75.77247184679887 |
| October | 84.56488091994801 | 18.160272196637678 |
| November | 10.933960634505228 | 14.499738918655652 |
June
July
August
September
October
November
Fig. A6 Chl-a-specific daily nitrate uptake rate (a) and daily phosphate uptake rate (b) of the phytoplankton community after NP enrichment experiment. Note each uptake rates were calculated from the nutrients concentrations at the start and the end of the NP treatments. Error bars denote mean SD
